# A non-catalytic function of a disintegrin and metalloprotease 10 determines hepatic progenitor cell fate

**DOI:** 10.1101/2024.08.14.607697

**Authors:** Birte Wöhner, Miryam Müller, Michael Pearen, Julia Köhn-Gaone, Luise Gorki, Rebecca Philippsen, Jully Gogoi-Tiwari, Clara John, Ludger Scheja, Freia Krause, Birgit Halwachs, Roja Barikbin, Dorothee Schwinge, Stefan Rose-John, Paul Saftig, Christoph Schramm, John K. Olynyk, Jörg Heeren, Janina EE Tirnitz-Parker, Grant A Ramm, Dirk Schmidt-Arras

## Abstract

During chronic liver disease, hepatocytes may undergo proliferative arrest, leading to the activation, expansion and differentiation of hepatic progenitor cells (HPCs). Here we observe that expression of A Disintegrin And Metalloprotease (ADAM) 10 is increased in human and murine chronic liver disease correlating with HPC expansion. We report that proteolytic processing of ADAM10 by ADAM9 and generation of an ADAM10 intracellular domain that translocates to the nucleus, rather than ADAM10 enzymatic activity is essential for the regulation of HPC gene expression and differentiation. Genetic loss of ADAM10 *in vitro* and *in vivo* enhances stemness gene expression, increases the accumulation of undifferentiated HPCs and promotes the formation of liver fibrosis. Taken together, we demonstrate that a non-catalytic function of ADAM10 is an essential regulator of HPC fate and HPC-driven regeneration. Our data ascribe a non-proteolytic function to ADAM proteases which may be a general concept in adult tissue stem cells.

## Introduction

The liver has an enormous capacity to regenerate. After acute damage, hepatocytes regenerate the lost tissue by proliferation and hypertrophy. However, hepatocyte proliferation is impaired under severe and chronic liver damage, resulting in a ductular reaction [1].

Ductular reactions are histological phenomena that encompass proliferation of ductular cells, inflammatory and extracellular matrix changes. They are thought to contain hepatic stem or progenitor cells (HPCs) that have the capacity to differentiate into hepatocytes or cholangiocytes, depending on the underlying injury stimulus, to aid hepatic repair and functional restoration [2, 3]. The cellular origin of HPCs has not been conclusively defined and may be highly context-specific [4]. There are several reports indicating that HPCs either reside in a stem cell niche close to or originate from biliary epithelial cells (BEC). This is further supported by the fact that both HPCs and BECs express common markers such as cytokeratin (CK) 19 and A6 antigen [5]. Recently, lineage-tracing experiments have demonstrated that under conditions of hepatic proliferative arrest, ductular cells give rise to newly formed hepatocytes *in vivo* [2].

In humans, activation of HPCs is observed in the vast majority of chronic liver diseases (CLD). Importantly, the degree of HPC proliferation positively correlates with disease severity [6, 7]. The associated mortality and morbidity have been linked to uncontrolled hepatic wound healing which progresses to liver cirrhosis and hepatocellular carcinoma (HCC) in severe cases [8]. During these processes the regenerative niche is comprised of inflammatory cells, activated hepatic stellate cells (HSCs) and HPCs that are in close spatial proximity and often in direct cell-cell contact [9, 10] . In this niche, myeloid cells secrete the tumor necrosis factor (TNF) family member TNF-like weak inducer of apoptosis (TWEAK) which was identified as a key HPC mitogen, signaling via its cognate receptor fibroblast growth factor-inducible 14 (Fn14/TNFRSF12A) [11]. Aberrant and prolonged activation of HPCs has been associated with progressive liver fibrosis and cirrhosis [10]. Furthermore, in HCC as well as cholangiocarcinoma, cells with a stem or progenitor cell phenotype can be identified and have been proposed to represent potential cellular precursors of these cancer types [8, 12, 13].

Cellular processes involved in progressive liver disease are regulated through complex molecular signaling networks. Proteolytic release of transmembrane ectodomains, also termed ectodomain shedding, is an irreversible post-translational mechanism to regulate protein function. Members of the A Disintegrin and Metalloprotease (ADAM) family are key mediators of ectodomain shedding. The family member ADAM10 was initially identified as a major sheddase for Notch receptors, thereby enabling Notch signaling [14]. As such, ADAM10 is involved in organogenesis and the maintenance of adult tissue stem cells [15]. We recently observed that while ADAM10 is dispensable for liver development, it is essential for the maintenance of adult liver homeostasis. Genetic loss of ADAM10 in hepatocytes, cholangiocytes and HPCs resulted in spontaneous hepatocyte necrosis and concomitant liver fibrosis, which was linked to aberrant bile acid transporter expression. In these mice an enhanced expansion of HPCs was seen in the absence of ADAM10, suggesting that ADAM10 mediates fundamental aspects of HPC biology [16].

In order to address the role of ADAM10 in HPCs in more detail, we aimed to analyze the effects of genetic ADAM10-deficiency in HPCs *in vitro* and *in vivo*. We now demonstrate that under conditions of chronic liver damage *Adam10* expression is significantly up-regulated and correlated with HPC expansion in mice and humans. Proteolytic processing of ADAM10 by ADAM9 generates an ADAM10 intracellular domain (ICD) within HPCs. Importantly, the nuclear translocation of ADAM10 ICD, and not ADAM10 proteolytic activity, restricts HPC expansion and primes HPCs to hepatocyte differentiation through enhanced shuttling of hepatocyte nuclear factor (HNF) 4α to the nucleus and repression of stemness genes. Accordingly, genetic loss of ADAM10 *in vivo* leads to enhanced HPC expansion, an increased expression of stem cell-associated marker genes and augmented liver fibrosis. Taken together, we identified a non-catalytic function of ADAM10 in HPCs, which is critically required for HPC-mediated liver regeneration.

## Material and Methods

### Animal models

All mice used in this study were on a C57BL/6N or a C57BL/6J background. ADAM10^fl/fl^ mice [17] were bred to ROSA26-STOP-tdTomato and CK19-CreERT2 mice.

All mice were housed under controlled conditions (specified pathogen free, 22°C, 12-hour day-night cycle) and fed a standard laboratory chow *ad libitum*. Animals received ethical care, according to the criteria outlined by the government of Germany and the European Union.

In the thioacetamide (TAA) model, mice were supplied with water containing 300 mg/L TAA (Sigma-Aldrich) for up to 96 days. TAA-containing water was replaced every second day.

In the choline-deficient, ethionine-supplemented (CDE) model, mice were fed *ad libitum* a choline-deficient chow (MP Biomedicals, NSW, Australia) and water containing 0.15% DL-ethionine (Sigma-Aldrich) for up to 96 days.

All animal experiments were performed at Christian-Albrechts-University Kiel, Germany according to the criteria outlined by the government of Germany (V244-2170/2017(31-3/17)) or at Curtin University Perth, Australia in accordance with the Australian code for the care and use of animals for scientific purposes, with local animal ethics committee approval.

### Human study objects

Liver samples from primary sclerosing cholangitis (PSC) patients were collected at University Medical Center Hamburg-Eppendorf as described [18]. Liver samples from patients with type 2 diabetes and non-diabetic controls were obtained at the Department of General Visceral Surgery at University Hospital Ulm, as described [19]. All patients gave written informed consent, and the studies were approved by the local ethics committee (permission numbers PV4081, Hamburg; 112/2003, Ulm) and the South Metropolitan Health Service Human Research Ethics Committee (protocol 96/37).

### Histological analysis and molecular techniques

see Supplementary Experimental Procedure

### Statistics

Microarray data processed with GEO2R used Linear Models for Microarray Analysis with Benjamini & Hochberg multiple testing correction.

Data are represented as mean ± s.e.m. Normal distribution of all experimental data was tested using Kruskal-Wallis. Comparisons between two groups were performed by applying Student *t* test and if not normally distributed using Mann-Whitney U test. Comparisons between multiple groups were performed by applying one-way ANOVA or if samples were not normally distributed with one-way ANOVA on ranks using Bonferroni’s post-hoc test. A *P* value < 0.05 was considered statistically significant.

## Results

### Adam10 expression is upregulated in chronic liver disease

Genetic deficiency of ADAM10 in hepatocytes, cholangiocytes and HPCs in mice results in spontaneous hepatocyte death and a concomitant accumulation of HPCs [16]. We hypothesized that ADAM10 is essential for the regulation of HPC biology and therefore assessed hepatic *Adam10* expression in murine and human CLD accompanied by ductular reaction and HPC expansion. Analysis of available transcriptomic data from patients suffering from alcoholic hepatitis compared to healthy controls [20] revealed a significant increase in *Adam10* expression under diseased conditions (Fig.1 a). *Adam10* expression correlated positively with transcript levels of *Fn14*, which is not detected in healthy liver and mainly expressed by HPCs activated during CLD (Fig.1 a). Also patients with type 2 diabetes (T2D) [19] or primary sclerosing cholangitis (PSC) [18] both displayed significantly elevated levels of hepatic *Adam10* expression (Fig.1 b). In particular, in liver biopsies of patients with PSC, ductular reactions with expansion of HPCs are generally observed [8]. Mice with a hepatocyte-specific nuclear factor kappa-B essential modulator (NEMO)-deficiency display spontaneous development of steatohepatitis and hepatocellular carcinoma [21]. Mdr2^-/-^ mice that are deficient for the ABC transporter B family member 4, involved in bile acid secretion, suffer from cholestatic damage, inflammation and fibrosis, accompanied with ductular reaction and HPC expansion [22]. Again, analysis of available transcriptomic data in NEMO-deficient mice [21] and Mdr2^-/-^ mice, revealed that *Adam10* expression in the liver was significantly increased and correlated to *Fn14* expression (Fig.1 c+d).

**Figure 1:**
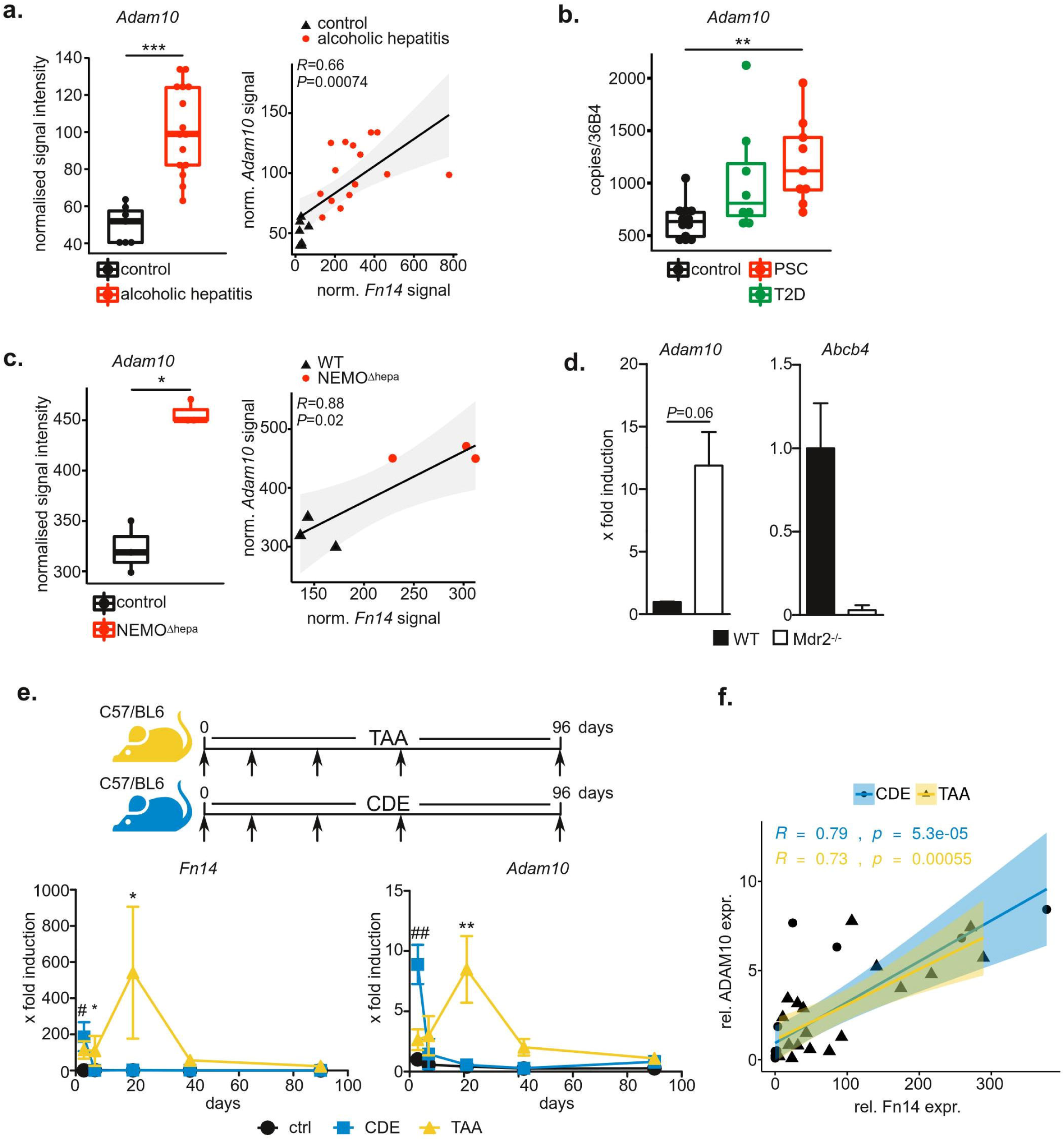
*Adam10* expression is upregulated in both, murine and human chronic liver disease and correlates with liver progenitor cell expansion as assessed by *Fn14* expression. **a.***Adam10* expression is elevated and correlates with *Fn14* expression in liver biopsies from patients with alcoholic hepatitis as assessed by microarray profiling [20]. **b.** ADAM10 expression is upregulated in patients with type 2 diabetes (T2D) or primary sclerosing cholangitis (PSC). **c.** Mice with liver-specific NEMO (IKKγ)-deficiency [21] display increased *Adam10* expression that correlates to *Fn14* expression in liver biopsies as assessed by microarray profiling. **d.** Expression of ADAM10 is upregulated in the cholestatic Mdr2^-/-^ model. *Adam10* and *Mdr2/Abcb4* expression was assessed by qRT-PCR. *Adam10* expression is upregulated only in female Mdr2^-/-^ mice which display a pronounced ductular reaction. **e.** C57BL/6 mice were subjected either to TAA in the drinking water or to a CDE diet. Kinetic changes in *Adam10* and *Tnfrsf12a/Fn14* expression was assessed by qRT-PCR on total liver RNA . **f.** Significant correlation of *Adam10* and *Fn14* expression in CDE- or TAA-treated mice indicates a prominent role of ADAM10 in HPC-mediated liver regeneration. Data are mean ± s.e.m. \**P*<0.05, \*\**P*<0.01, \*\*\**P*<0.001, Unpaired one-tailed Mann-Whitney U (a,c), Kruskal-Wallis with Bonferroni post-hoc test (b), One-way ANOVA (e), Pearson correlation (a, c, f), For microarray analysis, *P* values are adjusted for multiple testing correction.

In order to further assess the function of ADAM10 in CLDs, we subjected mice to two different CLD models (Fig.1 e). Choline-deficient ethionine-supplemented diet (CDE), as well as administration of thioacetamide (TAA) are two widely used mouse models to study hepatic steatosis, chronic inflammation, liver fibrosis and HPC expansion [9, 23]. We observed a transient increase in transcript levels for *Fn14*, which is closely associated with activation and expansion of HPCs [11] (Fig.1 e). *Adam10* expression was also transiently increased during CDE or TAA treatment (Fig.1 e) and significantly correlated with *Fn14* expression (Fig.1 f). Taken together, our data suggest that ADAM10 is actively involved in HPC-driven liver regeneration.

### Notch-dependent biliary differentiation of hepatic progenitor cells is independent of ADAM10

Differentiation of hepatoblasts as well as HPCs into the cholangiocytic lineage was previously linked to Notch2 activity [24]. ADAM10 is the predominant Notch α secretase in most tissues investigated [14, 16]. We therefore analyzed the capacity of the well-characterized murine HPC line BMOL [25] to form biliary tubes, which is a characteristic of functional cholangiocyte differentiation, after siRNA-mediated knock-down of ADAM10 (Fig.S1, S2 c) in a Notch2-dependent Matrigel assay [16] (Fig.S2 a). While the γ-secretase inhibitor DAPT reduced biliary tube formation, siRNA-mediated suppression of ADAM10 had no effect on branching or tube length of biliary tubes (Fig.S2 b), indicating that ADAM10 does not process Notch2 in HPCs. Accordingly, neither expression of the Notch ligand *Jag1*, nor the Notch target genes *Hes1* and *Hey2* were altered following knockdown of ADAM10 (Fig.S 2 c+d). Taken together, these data indicate that ADAM10 is dispensable for Notch signaling in HPCs.

### Loss of ADAM10 increases stemness of hepatic progenitor cells independent of its catalytic activity

In order to analyze the consequences of ADAM10-deficiency, we assessed expression of genes linked to HPC differentiation. Interestingly, we detected a substantial increase in both, albumin as well as *Krt19* (the gene encoding CK19) in ADAM10-deficient BMOL cells (Fig.2 a,b). This finding is reminiscent of *Alb* and *Krt19* co-expression in hepatoblasts during development [26]. While expression of the hepatocytic transcription factor *HNF4α* was not altered with *Adam10* knockdown, expression of *Hnf6*, normally expressed in cholangiocytes and HPCs, was significantly increased, as well as levels of the hepatic transcription factor Liver X receptor alpha (*Lxrα*) (Fig.2 c). In addition, expression of the stem cell markers *Sox9*, *Cd24a* and *Cd133* were significantly elevated following knockdown of *Adam10* (Fig.2 d). These findings were specific to ADAM10 knockdown, as we did not observe strong alterations in gene expression, when we knocked down the close homologue ADAM17 in BMOL cells (Fig.2 a-d). Assessment of protein levels by immunofluorescence, confirmed significant up-regulation of CD133 and CK19 in the absence of ADAM10 (Fig.2 e+f). Furthermore, we noticed that ADAM10-deficient BMOL cells displayed an altered, more rounded morphology with smaller nuclei as compared to control cells (Fig.2 e). Taken together, these data suggest that the absence of ADAM10 results in the transition of HPCs towards a non-lineage-committed, stem cell-like phenotype.

**Figure 2:**
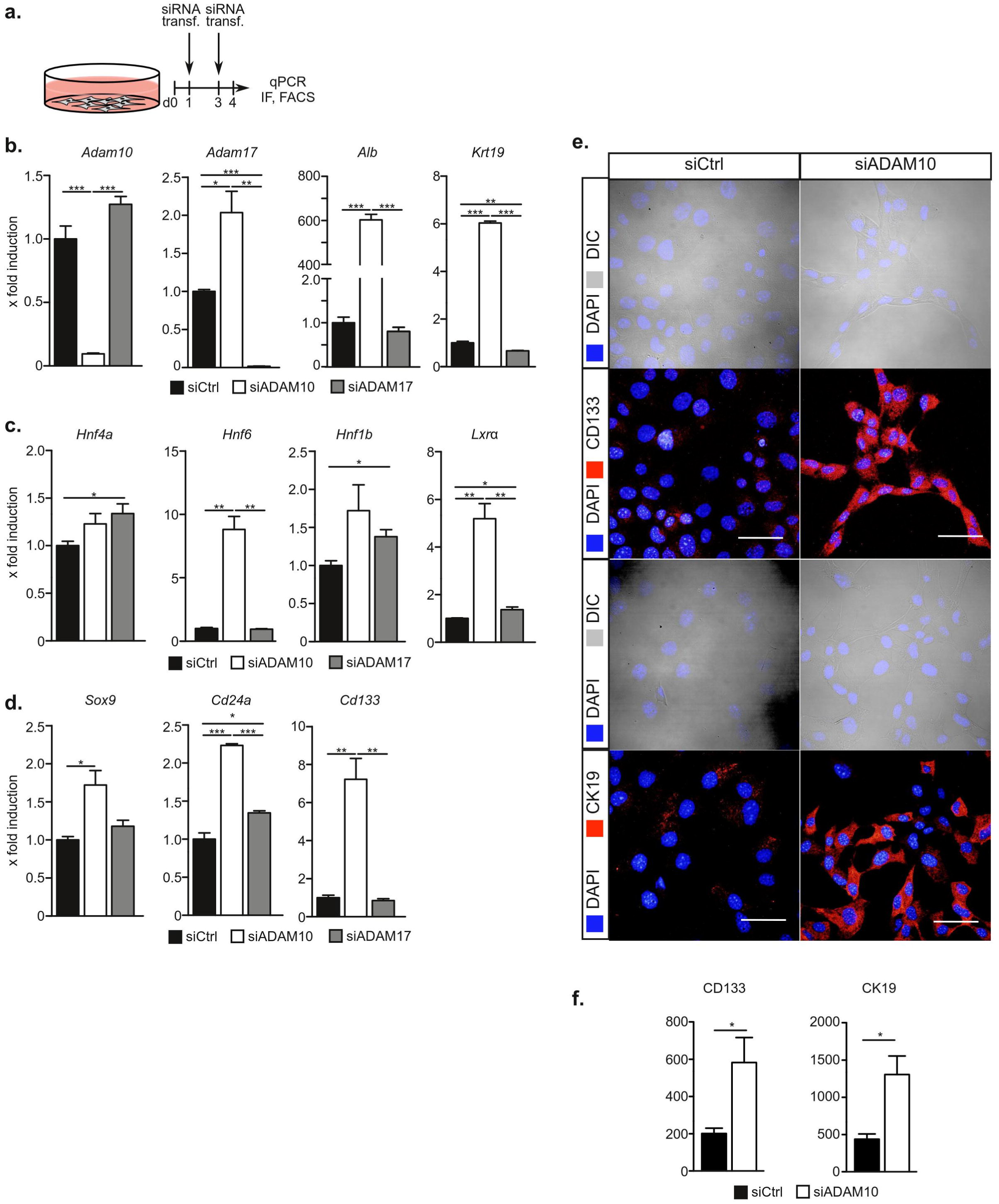
Expression of stemness-related genes is regulated by ADAM10. **a.** Experimental outline as performed in b-f. **b.** Expression of both, albumin and *Krt19* is up-regulated in the absence of ADAM10 but not ADAM17. **c.** Expression of the hepatic transcription factors *Hnf6* and *Lxrα* is increased in the absence of ADAM10 but not ADAM17. **d.** Expression of the stemness-associated transcription factor Sox9 and surface markers *Cd24a* and *Cd133* is upregulated in the absence of ADAM10 but not ADAM17. **e.** Morphological changes and increased expression of CD24α and CD133 in ADAM10-deficient HPCs as assessed by immunofluorescence on siRNA-depleted BMOL cells. **f.** Quantification of CK19 and CD133 immunofluorescent signals. Data are mean ± s.e.m. n=3 (b-d), n=4 (f) \**P*<0.05, \*\**P*<0.01, \*\*\**P*<0.001, unpaired, two-tailed Student’s t test (b-d), unpaired, two-tailed Mann-Whitney U (f).

Given the significant increase in *Lxrα* expression in the absence of ADAM10, we investigated whether ADAM10-dependent alterations in gene expression are linked to LXRα activity. We therefore employed the previously described inverse LXR agonist SR9243 that impairs LXR α activity [27]. While we observed impaired expression of LXRα target genes *Fasn* and *Scd1*, we did not observe alterations in either *Alb* or *Krt19* expression or in the expression of selected hepatic transcription factors or stem cell markers (Fig.S3 a-e) in the presence of SR9243. Thus, the LXR nuclear receptors do not account for ADAM10-dependent regulation of gene expression.

In order to determine whether proteolytic activity of ADAM10 is needed to regulate gene expression, we employed recombinant (r) tissue inhibitor of metalloproteinase (TIMP)-1, a physiological inhibitor of metalloproteases including ADAM10 [28], as well as the ADAM10 small molecule inhibitor GI254023X [29]. Both agents prevent ADAM10 catalytic activity by binding to the active site. Surprisingly, only siRNA-mediated downregulation of ADAM10 protein levels, but not inhibition of ADAM10 catalytic activity by rTIMP-1 or GI254023X led to a marked increase in *Alb* and *Krt19* expression (Fig.3 a+b), an upregulation of *Hnf6*, *Lxrα* (F ig.3 c), and the stem cell markers *Cd24a* and *Cd133* (Fig.3 d). A slight upregulation of *Hnf4a* and *Sox9* was seen after treatment with rTIMP-1 but not GI254023X, most likely resulting from rTIMP-1-mediated inhibition of other proteases than ADAM10. We therefore concluded that a non-catalytic function of ADAM10 regulates HPC gene expression and primes HPCs for differentiation.

**Figure 3:**
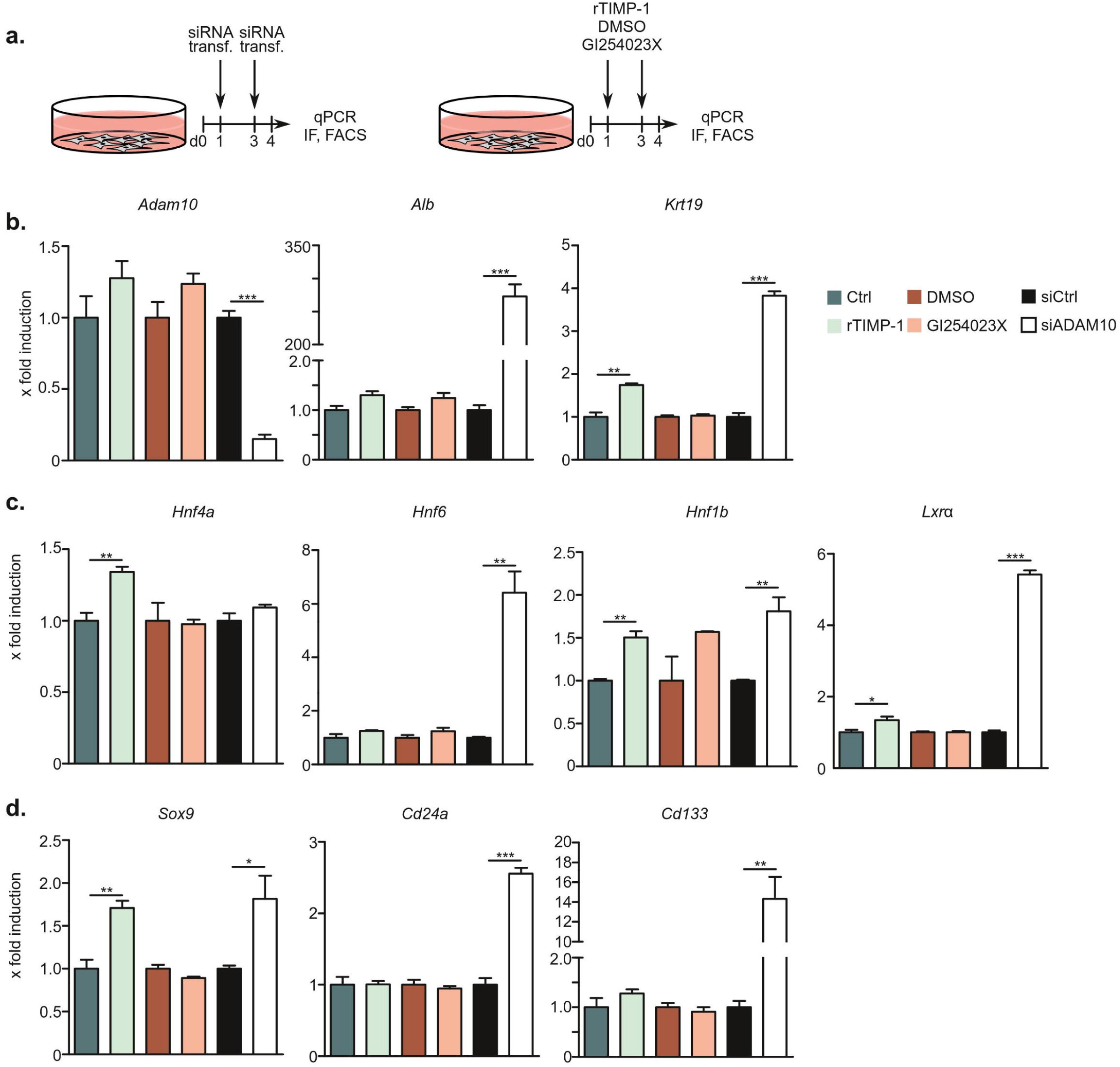
Enzymatic activity of ADAM10 is dispensable for transcriptional changes in HPCs. **a.** Experimental outline as performed in b-d. Expression of differentiation-related genes (**b**), hepatic transcription factors (**c**) or stemness-related genes (**d**) is altered in the absence of ADAM10 protein, but unaltered if ADAM10 catalytic activity is inhibited by recombinant TIMP-1 or the small molecule ADAM10 inhibitor GI254023X. Data are mean ± s.e.m. n=3, \**P*<0.05, \*\**P*<0.01, \*\*\**P*<0.001, unpaired, two-tailed Student’s t test (b-d).

### ADAM10 is processed by ADAM9 and translocates to the nucleus in hepatic progenitor cells

We hypothesized that proteolytic processing of ADAM10 might direct an ADAM10 ICD to the nucleus. We therefore assessed ADAM10 localization in BMOL cells by immunofluorescence using an antibody directed against the C-terminus of ADAM10. In control cells, ADAM10 localized to a perinuclear region as previously described [30], but substantial localization was also observed in the nucleus (Fig.4 a). Signals for ADAM10 were completely absent in cells with knockdown of ADAM10 (Fig.4 a). The effect was specific for ADAM10, as knockdown of ADAM17 did not alter nuclear localization of ADAM10 (Fig.S4 a). We observed similar results using an anti-Flag antibody in BMOL cells with CRISPR-mediated endogenous C-terminal Flag-tagging of ADAM10 (Fig.S4 b+c). Liver tissue sections from mice subjected to CDE diet or TAA treatment displayed nuclear ADAM10 staining in cells of the ductular reaction and hepatocytes but not other non-parenchymal cells, suggesting a role of nuclear ADAM10 during LPC-mediated regeneration (Fig.4 b). Nuclear localization of ADAM10 was also observed in the human HPC line HepaRG (Fig.S4 d+e) and in liver biopsies from patients with CLD associated with hepatitis C virus-infection (Fig.4 c). Here, nuclear ADAM10 was detectable in strings of ductular cells, presumably HPCs (Fig.4 c, red arrow heads) and in adjacent hepatocytes (Fig.4 c, black arrow heads), that may have arisen through the differentiation of HPCs.

**Figure 4:**
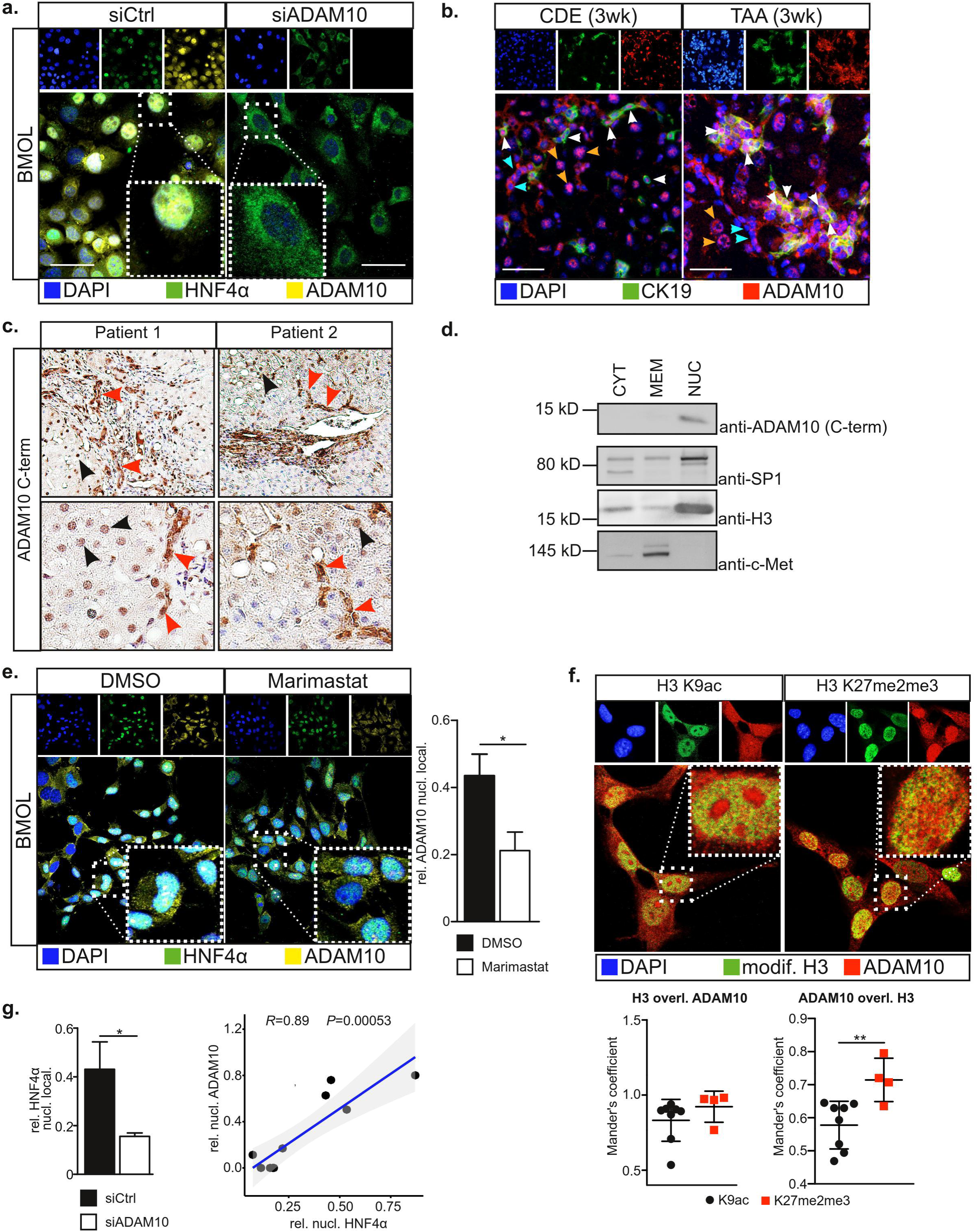
An ADAM10 intracellular domain (ICD) translocates to the nucleus of LPCs and correlates with HNF4α nuclear translocation. **a.** Nuclear translocation of HNF4α and ADAM10 ICD as assessed by immunofluorescence of siRNA-transfected BMOL cells using an antibody raised against ADAM10 C-terminus. **b.** ADAM10 localizes to the nucleus in cells of the ductular reaction (white arrow heads) and hepatocytes (yellow arrow heads), but not to other non-parenchymal cells (cyan arrow heads) in liver tissue sections of mice with chronic liver disease either 3 weeks after CDE diet or TAA treatment. CK19 labels cholangiocytes and liver progenitor cells. **c.** Liver biopsies from patients suffering from chronic liver disease, associated with hepatitis C virus-infection, display nuclear translocation of ADAM10 ICD in hepatic ductular reactions. Nuclear ADAM10 is in particular found in strings with expanding cells of the ductular reaction, presumably hepatic progenitor cells (red arrow heads), but also detected in hepatocytes (black arrow heads). **d.** A 10kD C-terminal fragment of ADAM10 translocates to the nucleus as determined by subcellular fractionation and immunoblotting. Purity of subcellular fractionation was assessed by immunoblotting against the transmembrane protein c-Met, nuclear histone H3 and the transcription factor SP1. CYT: cytoplasm, MEM: membrane, NUC: nucleus **e.** Nuclear localization of ADAM10 is significantly reduced in the presence of the broad spectrum metalloprotease inhibitor Marimastat. **f.** Nuclear ADAM10 localizes to histone H3 methylated at Lys-27 (H3 K27me2me3; transcriptionally inactive) rather than to H3 acetylated at Lys-9 (H3 K9ac; transcriptionally active) in BMOL cells. **g.** HNF4α nuclear translocation is significantly decreased in the absence of ADAM10. The extent of HNF4α nuclear shuttling correlates with ADAM10 ICD nuclear localization. Data are mean ± s.e.m. n=8-9 (b), n=4 (e), n=4-8 (f) \**P*<0.05, \*\**P*<0.01, one-tailed unpaired Student’s t-test (e), one-tailed unpaired Mann-Whitney U (f, g), Pearson correlation (g).

Multiple sequence alignment (MSA) revealed high conservation of ADAM10 ICD between different vertebrate species (Fig.S5 a). While we did not detect a classic nuclear localization sequence using PredictNLS [31], MSA-based prediction of murine ADAM10 ICD (Fig.S5 b) using LocTree3 [32] confirmed potential nuclear localization of ADAM10 ICD with high expected accuracy (Fig.S5 c). Indeed, we detected an approximately 10 kDa-sized ADAM10 ICD in BMOL cells (Fig.S5 d), that was exclusively detected in the nuclear fraction when cells were subjected to subcellular fractionation (Fig.4 d). In accordance with proteolytic processing by metalloproteases, nuclear ADAM10 was significantly reduced, when BMOL cells were treated with the broad-spectrum metalloprotease inhibitor Marimastat (Fig.4 e). In the nucleus, ADAM10 ICD was co-localized with Histone H3 di- or trimethylated at Lys-27 (H3K27me2me3), rather than with H3 acetylated at Lys-9 (H3K9ac) (Fig.4 f). This indicates that ADAM10 ICD localizes to transcriptionally inactive chromatin. Interestingly, in both, BMOL and HepaRG cells, nuclear localization of the transcription factor HNF4α correlated with the nuclear localization of ADAM10 (Fig.4 a+g and Fig.S4 d+e). This correlation was also observed in Marimastat-treated cells (Fig.S4 e). In the absence of ADAM10, HNF4α was predominantly localized to the cytoplasm (Fig.4 a+g). In line with these observations, upregulation of *Krt19* and *Cd133* expression was inversely correlated with nuclear localization of ADAM10 and HNF4α (Fig.S6 a-c).These data suggest that ADAM10 ICD is needed for the nuclear shuttling of HNF4α.

ADAM10 can be proteolytically processed by ADAM9 and ADAM15 (Fig.S7 a) [33]. ADAM9 expression was readily detectable by immunofluorescence in BMOL cells (Fig.S7 b). siRNA-mediated downregulation of ADAM9 or ADAM15 impaired ADAM10 ICD formation (Fig.S7 c+d) and significantly lowered ADAM10 nuclear localization (Fig.5 a, b). In line with the assumption that ADAM10 ICD regulates HNF4α shuttling, we observed that in ADAM9- or ADAM15-depleted cells nuclear localization of HNF4α was significantly lowered (Fig.5 a). In addition, downregulation of ADAM9 increased *Alb* expression, presumably via proteolytic processing of ADAM10, however to a lower extent than depletion of ADAM10 itself (Fig.5 c). Accordingly, increased expression of *Hnf6* and *Lxrα,* as well as stem cell-related genes *Sox9, Cd24a* and *Cd133* (Fig.5 c) was detectable in the absence of ADAM9. In agreement with a transcriptional repressive role of ADAM10 ICD, we detected decreased levels of Lys-27 trimethylated histone H3 (H3K27me3) in the absence of ADAM9 or 10 (Fig. 5 d). In line with a proposed biological role of ADAM9 and 15 in HPCs *in vivo*, we observed that expression of *Adam9* and *Adam15* were significantly increased in CDE- or TAA-treated mice (Fig.5 e) and displayed similar expression patterns as transcripts of *Fn14* and *Adam10* (Fig. 1 e). Supporting an association with CLD, *Adam9* and *Adam15* expression were significantly increased in liver biopsies from patients with alcoholic hepatitis [20] (Fig.5 f) and *Adam15* expression was significantly elevated in livers of hepatocyte-specific NEMO (IKKγ)-deficient mice [21] (Fig.5 g).

**Figure 5:**
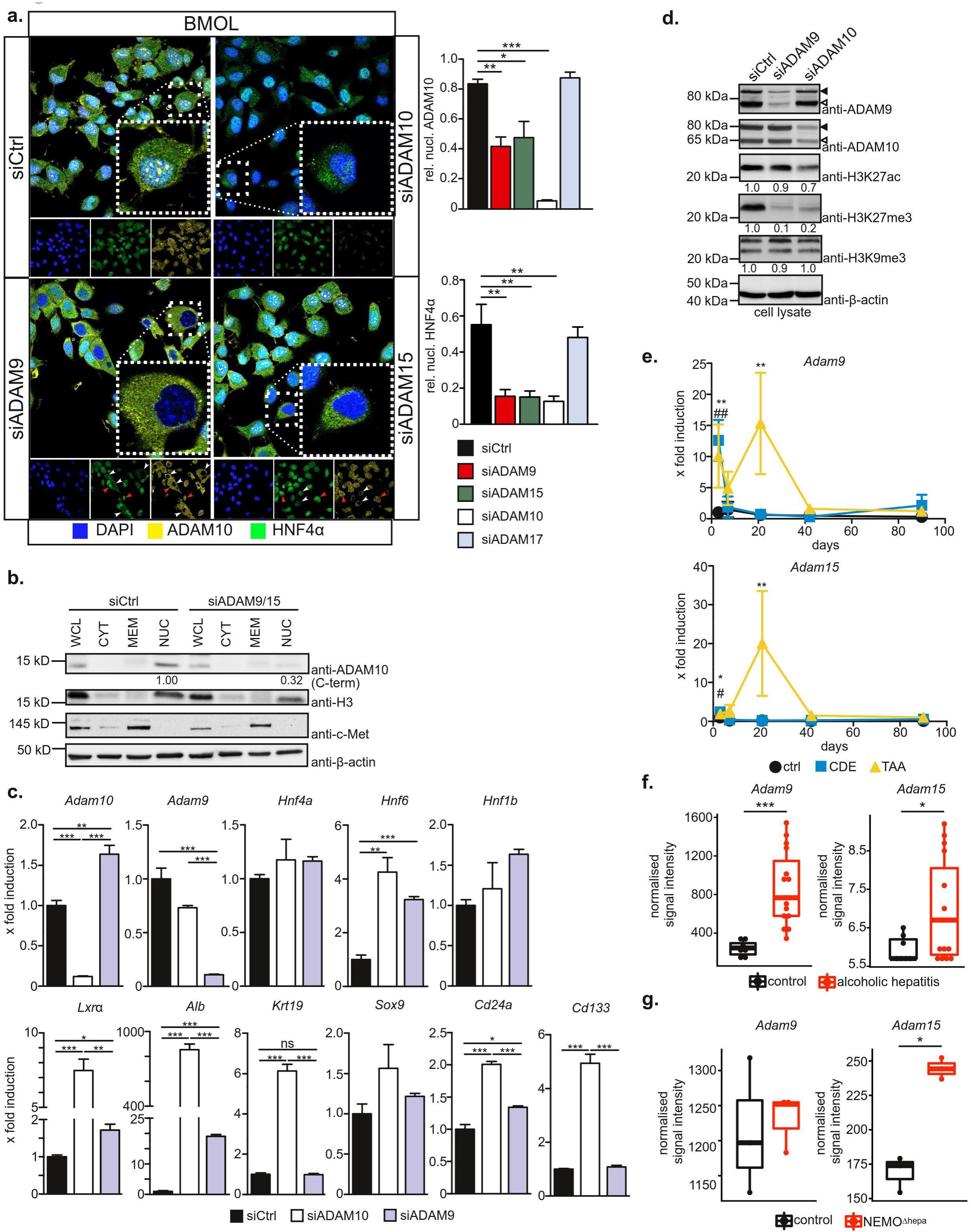
Nuclear localization of ADAM10 in HPCs depends on its processing by ADAM9 or ADAM15. **a.** siRNA-mediated suppression of ADAM9 or ADAM15 significantly lowers nuclear localization of ADAM10 ICD and the transcription factor HNF4α. **b.** siRNA-mediated concomitant suppression of ADAM9 and ADAM15 strongly reduces nuclear localization of ADAM10 ICD as assessed by subcellular fractionation and immunoblotting against ADAM10. Purity of fractionation was assessed by immunoblotting against the transmembrane protein c-Met and the nuclear histone H3. WCL: whole cell lysate, CYT: cytoplasm, MEM: membrane, NUC: nucleus **c.** siRNA-mediated suppression of ADAM9 induces similar changes in HPC gene expression as ADAM10-deficiency. Differential expression of the indicated genes was assessed by qRT-PCR on total liver RNA. **d.** Histone H3K27 trimethylation is lost upon siRNA-mediated suppression of ADAM9 or ADAM10. BMOL cells were treated with the indicated siRNAs and total cell lysates were analyzed by immunoblotting using the indicated antibodies. **e.** Expression of ADAM9 and ADAM15 is increased during the course of the CDE and TAA murine models of chronic liver disease. The expression of ADAM9 and ADAM15 follows ADAM10 and Fn14 expression (see Figure 1 e). Differential expression of the indicated genes was assessed by qRT-PCR. **f.** *Adam9* and *Adam15* expression is elevated in liver biopsies of alcoholic hepatitis patients [20] as assessed by microarray profiling. **g.** Loss of hepatic NEMO (IKKγ) [21] induces *Adam15*-expression in the liver as analyzed by microarray profiling. Data are mean ± s.e.m. n=4-5 (a), n=3 (c) \**P*<0.05, \*\**P*<0.01, \*\*\**P*<0.001, Unpaired, two-tailed Student’s t-test (f, *Adam9*), unpaired one-tailed Student’s test (a) unpaired one-tailed Mann-Whitney U (f, *Adam15, g*), One-way ANOVA (e), One-way ANOVA on ranks with Bonferroni’s post-hoc test (a, c). For microarray analysis, *P* values are adjusted for multiple testing correction.

Taken together, these data suggest that in HPCs ADAM10 is proteolytically processed by ADAM9/15 to produce an ADAM10 ICD that can translocate to the nucleus, assist in HNF4α nuclear shuttling and the transcriptional inactivation of stem cell genes in order to regulate HPC biology and permit HPC differentiation.

### Accumulation of undifferentiated hepatic progenitor cells in the absence of ADAM10 in vivo

In order to assess if differentiation of HPCs is dependent on ADAM10 *in vivo*, we generated mice with ADAM10-deficiency in CK19^+^ cells comprising biliary epithelial cells (BECs) and HPCs (ADAM10^iΔHPC/BEC^) by breeding ADAM10^fl/fl^::R26^tdTomato^ mice to mice with tamoxifen-inducible CreERT2 under the control of the *Krt19* (CK19) promoter [34] . As a control strain (ADAM10^WT^), R26^tdTomato^ mice were bred to CK19-CreERT^2^ mice (Fig. 6 a). In both mouse lines, induction of Cre activity is accompanied by expression of a tdTomato reporter allele. We subjected these mice to the TAA chronic liver injury and fibrosis model (Fig.6 b). Alterations in relative body weight were similar in ADAM10-deficient and -proficient mice (Fig. 6 c). We also did not observe large differences in relative liver and spleen weight in ADAM10^iΔHPC/BEC^ mice (Fig.6 d). A6^+^ cells, comprising HPCs and BECs, displayed recombined alleles as assessed by tdTomato expression (Fig.6 e), which was enhanced when mice were challenged with TAA (Fig. 6 e). Loss of ADAM10 in HPC/BECs led to a significantly enhanced proportion of recombined alleles in A6^+^ cells (Fig. 6 e). As TAM-treated ADAM10^WT^ mice reacted similar compared to vehicle-treated ADAM10^iΔHPC/BEC^ mice, we continued our experiments with ADAM10^iΔHPC/BEC^ mice only.

**Figure 6:**
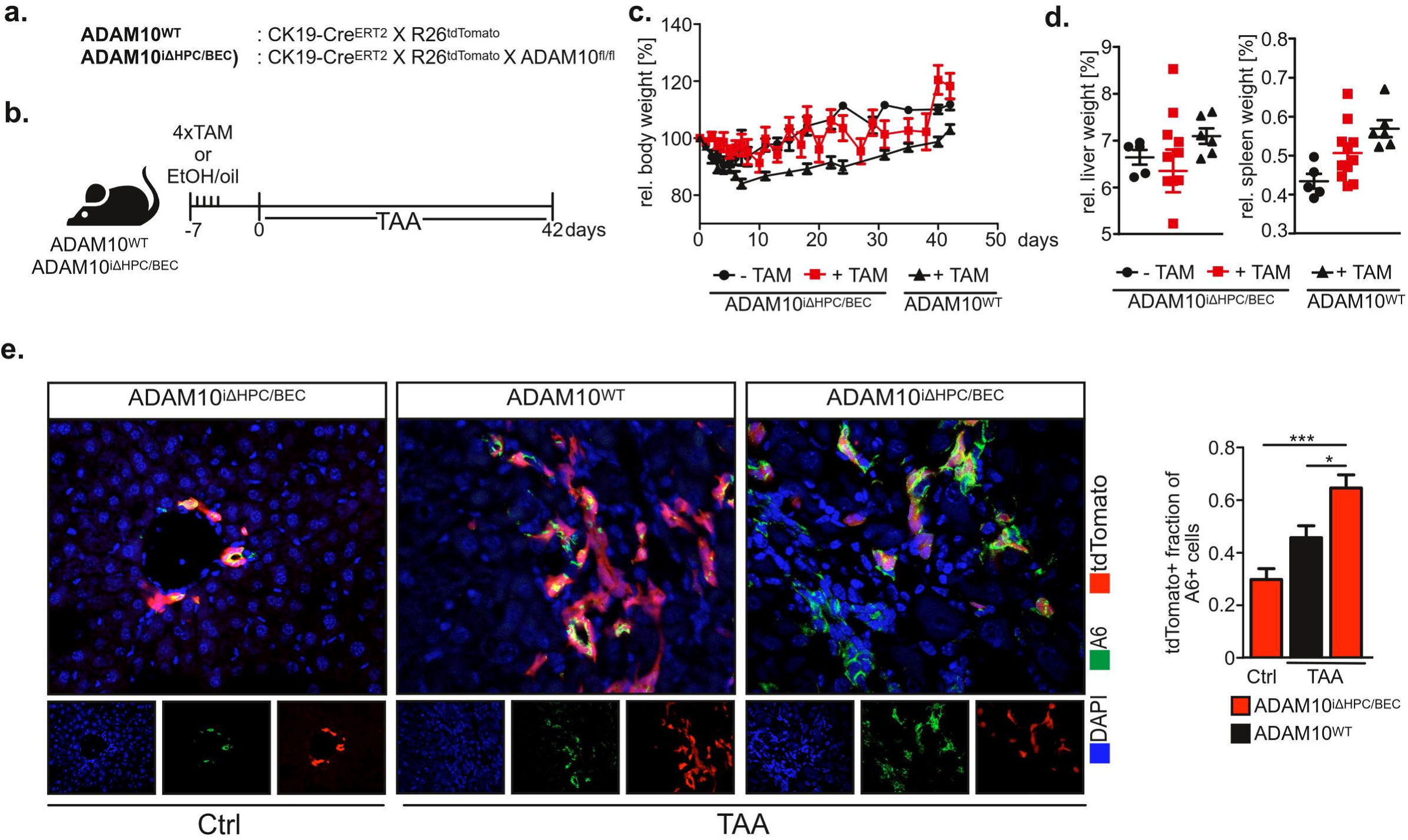
A mouse model of ADAM10 genetic deficiency in hepatic progenitor cells. **a.** Genetic mouse models used in Figure 6-8. Cre activity induces expression of a tdTomato reporter in CK19^+^ cholangiocytes and hepatic progenitor cells in both mouse lines. **b.** Experimental outline as performed in Figure 6-8. Tamoxifen (TAM)-induced Cre activity and subsequent loss of ADAM10 expression in CK19^+^ cells was induced by four consecutive tamoxifen injections. Subsequently mice were either left untreated or subjected to a thioacetamide (TAA) liver damage model. **c.** Body weight change during TAA model is similar in all three groups. **d.** Liver and spleen weight relative to body weight are not largely altered in the absence of ADAM10 in HPCs and cholangiocytes. **e.** Recombination efficiency in A6^+^ cells as assessed by tdTomato and A6 colocalization. Data are mean ± s.e.m. n=4-9 mice/group (c,d), n=3-4 mice/group with 4-5 images/sample analyzed (e), \**P*<0.05, \*\*\**P*<0.01, One-way ANOVA with Tukey’s post-hoc test (e).

In unchallenged TAM-treated ADAM10^iΔHPC/BEC^ mice, i.e. in the absence of parenchymal damage, we observed a small but significant increase in A6^+^ cells in ADAM10-deficient mice (Fig. 7 a). When mice were subjected to chronic liver damage induced by TAA, the number of A6^+^ cells was markedly increased in ADAM10-deficient cells (Fig. 7 a), indicating increased activation and proliferation of HPCs or an accumulation of HPCs due to impaired differentiation. In line with our *in vitro* experiments, expression of stemness-associated CD133 was significantly increased in HPCs with excised *Adam10* allele, as assessed by tdTomato expression in CD133^+^ cells (Fig. 7 b). Accordingly, we observed a significant increase in *Krt19*, *Cd133* and *Sox9* expression in total liver RNA in ADAM10^iΔHPC/BEC^ mice, which was not observed in control mice (Fig. 7 c and Fig.S8). The increase in stem cell markers in ADAM10-deficient HPCs was not linked to altered levels of known HPC mitogens including hepatocyte growth factor (*Hgf*), lymphotoxin β (*Ltb*) or TWEAK (*Tnf2f12*) (Fig. 7 c), further supporting the hypothesis of a cell-intrinsic effect of ADAM10 on HPC gene expression and cell fate.

**Figure 7:**
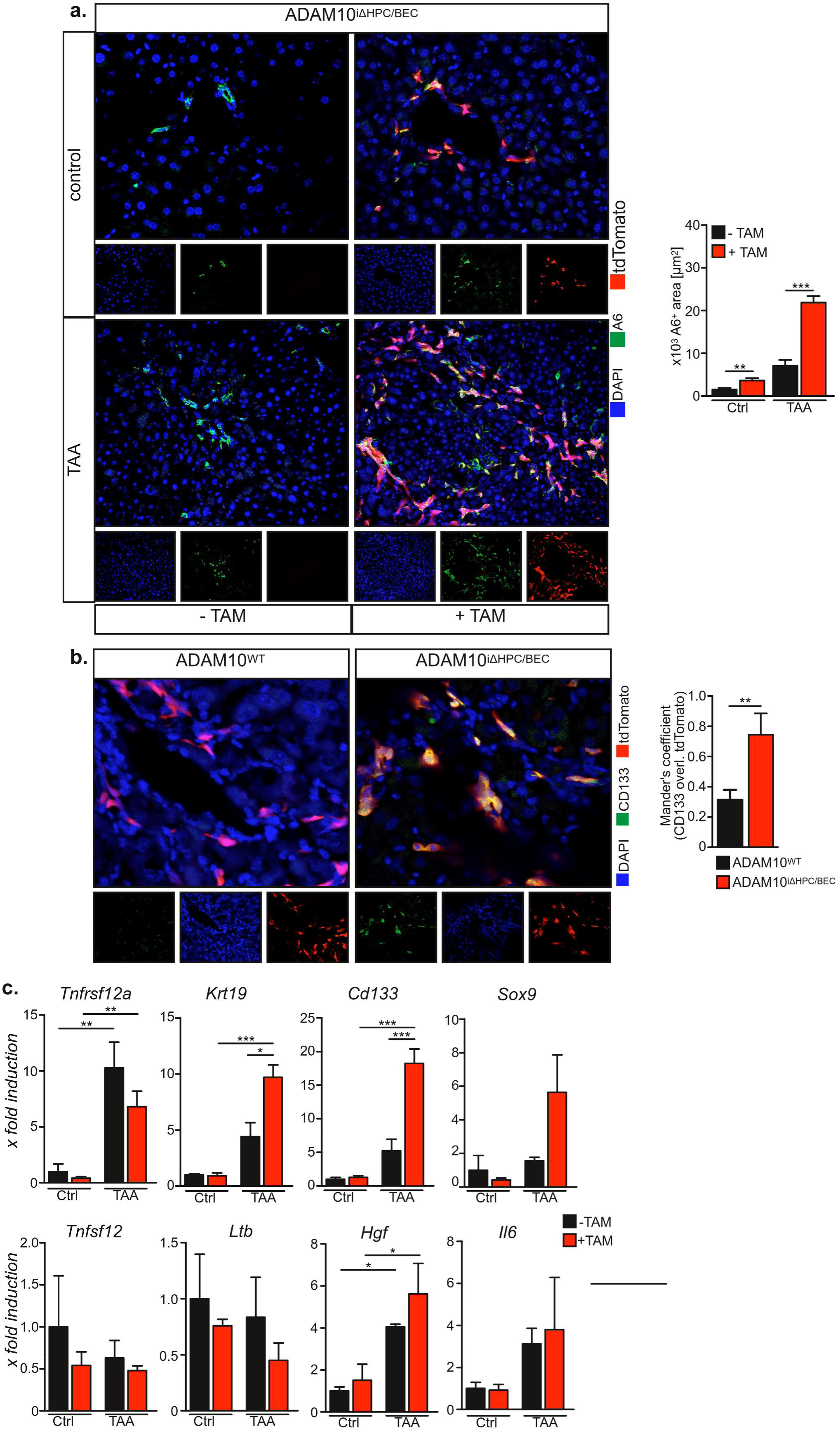
ADAM10 regulates hepatic progenitor cell fate *in vivo.* **a.** The number of A6^+^ cells is increased in the absence of ADAM10 in HPCs, both in unchallenged and TAA-treated mice. **b.** Loss of ADAM10 in HPCs increases proportion of CD133 in HPCs as assessed by tdTomato and CD133 colocalization. **c.** Expression of CD133, CK19/Krt19 and Sox9 is increased in livers with ADAM10-deficient HPCs, while expression of HPC mitogens like TWEAK/Tnfsf12, HGF, lymphotoxin β/LTb or IL-6 are similar in ADAM10-proficient or - deficient mice. Data are mean ± s.e.m. n=3 mice/group (a-c) with 4-5 images/sample analyzed (a-b) \**P*<0.05, \*\**P*<0.01, \*\*\**P*<0.001, unpaired two-tailed Student’s t-test (a), unpaired Student’s t-test with Welch’s correction (b), One-way ANOVA on ranks with Bonferroni’s post-hoc test (c).

Expansion of HPCs in chronic liver disease is linked to hepatic stellate cell (HSC) activation and liver fibrosis [9, 10]. We therefore assessed collagen deposition by Sirius Red staining of liver tissue sections. We observed a significant increase in collagen deposition in unchallenged ADAM10-deficient mice (Fig. 8 a+b) which correlated with the expansion of A6^+^ HPCs (Fig. 7 a). When subjected to TAA treatment, collagen deposition increased in both groups of mice, with Sirius Red-positive areas significantly elevated in ADAM10^iΔHPC/BEC^ mice (Fig. 8 b). Interestingly, while we observed an increase in fibrosis-associated gene expression upon TAA treatment, no significant differences between ADAM10-proficient and ADAM10-deficient animals was observed (Fig. 8 c). These findings suggest that hepatic stellate cell activation by undifferentiated HPCs is not linked to paracrine signals but rather to direct cell-cell contacts of the two cell types as has previously been proposed [35, 36]. While activated HSCs were barely detectable in control animals, we found increased numbers of αSMA^+^ HSCs in close proximity to HPCs within ADAM10-deficient livers (Fig. 8 d). Furthermore, the number of F4/80^+^ myeloid cells in close proximity to HPCs was elevated in ADAM10^iΔHPC/BEC^ mice as compared to control animals (Fig. 8 e).

**Figure 8:**
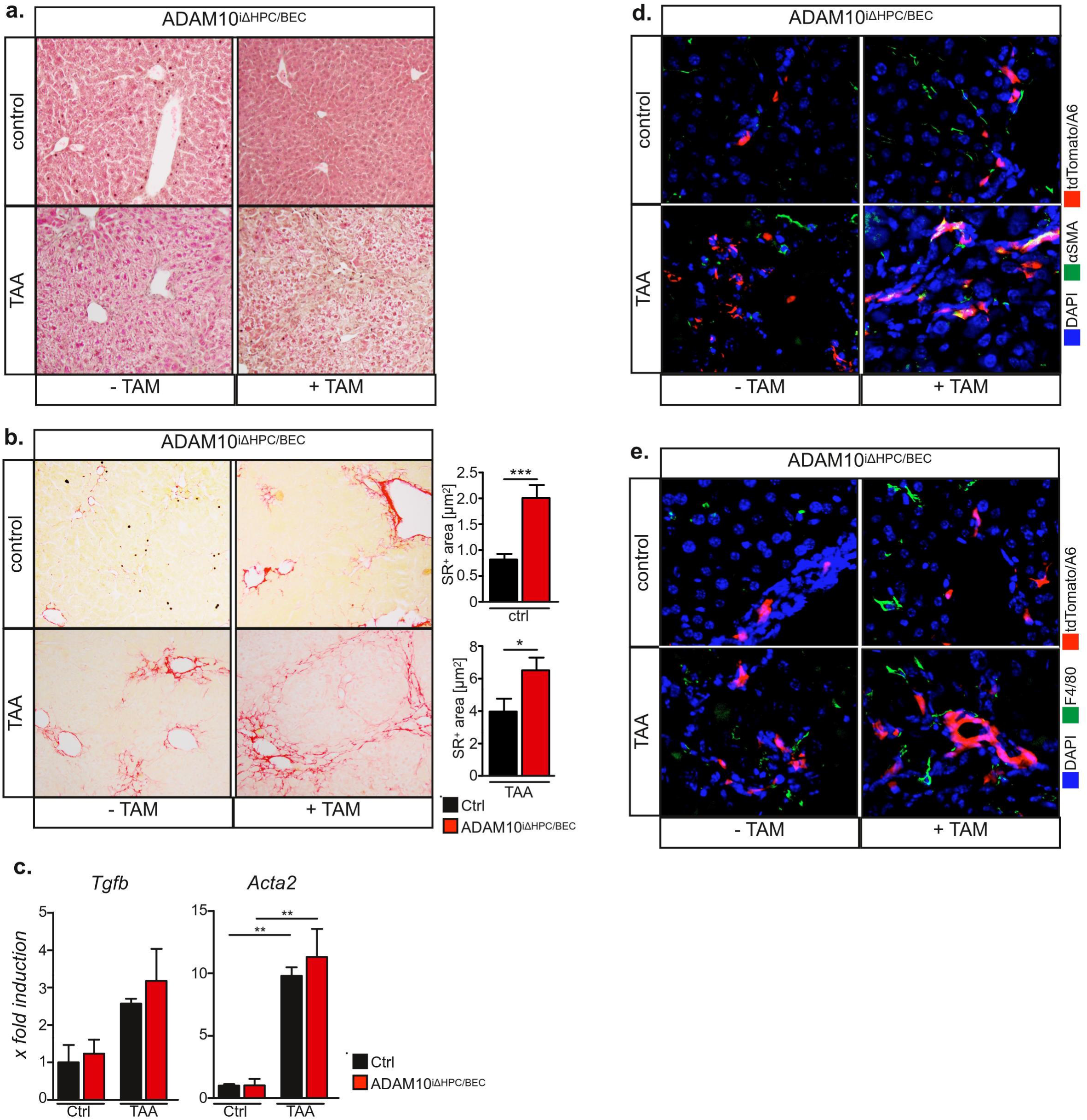
Loss of ADAM10 in HPCs aggravates liver fibrosis. **a.** H/E histochemistry of liver sections from untreated and TAA-treated animals. **b.** Increased collagen deposition in ADAM10^iΔHPC/BEC^ mice as assessed by Sirius Red histochemistry of liver sections from untreated (control) and TAA-treated animals. **c.** qRT-PCR analysis of fibrosis-associated genes on RNA isolated from untreated and TAA-treated animals. **d.** The number of activated αSMA+ hepatic stellate cells (HSCs) in close proximity to HPCs is increased in mice with ADAM10-deficient HPCs. HPCs in liver tissue sections were identified either by tdTomato fluorescence (ADAM10^iΔHPC/BEC^) or by A6 staining (control). HSCs were identified with anti-αSMA antibodies. **e.** The number of F4/80+ myeloid cells in close proximity to HPCs is increased in mice with ADAM10-deficient HPCs. HPCs in liver tissue sections were identified either by tdTomato fluorescence (ADAM10^iΔHPC/BEC^) or by A6 staining (control). Data are mean ± s.e.m. n=3 mice/group (b, c) with 4-5 images/sample analyzed (c) \**P*<0.05, \*\**P*<0.01, \*\*\**P*<0.001, Two-tailed unpaired Mann-Whitney U (b), one-way ANOVA on ranks with Bonferroni’s post-hoc test (c).

In summary, we demonstrate that loss of ADAM10 in HPCs *in vivo* impairs HPC differentiation resulting in prolonged HSC activation and augmented liver fibrosis.

## Discussion

HCC is one of the most frequently occurring cancers worldwide and the major cause of death in liver cirrhosis patients in western countries. CLD such as chronic viral hepatitis, metabolic liver disease, chronic alcoholic and non-alcoholic steatohepatitis predispose to cancer formation [37]. HPC proliferation is observed in all CLD and its amplitude strongly correlates with the underlying disease severity [6]. Due to the limitations in tracing this very heterogenous population, it is not clear whether they represent tumor-initiating cells or regulatory cells that may indirectly influence cancer outcome by regulating other hepatic cell types in the regeneration niche [8, 12, 13]. Furthermore, a functional relationship between HPCs and HSCs in the generation of liver fibrosis has been discussed [10]. While HPC mitogens such as TWEAK [11] and HGF [5] are known, there is little understanding of the mechanisms that regulate HPC differentiation.

In the present study, we analyzed the expression of the membrane-bound metalloprotease ADAM10 using human transcriptomic data. ADAM10 expression was elevated in human CLD such as alcoholic hepatitis, PSC and type 2 diabetes. We validated these findings using available murine transcriptomic data in both the CDE and TAA murine models of CLD and the Mdr2^-/-^ model of PSC. ADAM10 was also elevated in NEMO-deficient mice that develop HCC on the background of spontaneous steatohepatitis.

We previously generated mice with a liver specific knockout of ADAM10 using Alfp-Cre mice [16]. In these mice, genetic deficiency of ADAM10 started from day E9.5 on and therefore ADAM10 was lost in hepatoblasts and consequently absent in hepatocytes, cholangiocytes and HPCs. These mice displayed spontaneous hepatocyte necrosis with subsequent development of liver fibrosis. We also observed an accumulation of HPCs in these mice. However, this study did not determine whether HPC expansion was a consequence of hepatocyte death or linked to ADAM10-loss in HPCs [16] .

Here we demonstrate that genetic deficiency of ADAM10 in HPCs impairs the potential of HPCs to differentiate *in vitro* and *in vivo*. Mice with ADAM10-deficient HPCs displayed increased numbers of HPCs under control and diseased conditions. This was not due to increased expression of HPC mitogens. Rather, we observed that ADAM10-deficient HPCs expressed elevated levels of stem cell genes such as *Sox9*, *Cd24a* and *Cd133*, suggesting that ADAM10 deletion impaired HPC differentiation, resulting in the detection of increased numbers of HPCs. The elevation in HPC numbers in ADAM10^iΔHPC/BEC^ mice correlated with increased numbers of activated HSCs and elevated collagen as well as increased detection of F4/80^+^ cells, which include recruited monocyte-derived macrophages and liver-resident Kupffer cells. These data suggest that HPC proliferation within the ductular reaction, fibrogenesis and inflammation are tightly co-regulated, as demonstrated in hepatitis C virus-infected patients before and after liver transplantation [38].

Interestingly, the increase in HPC stemness was not linked to an absence of ADAM10 enzymatic activity. Instead we observed nuclear localization of an ADAM10 ICD that co-localized with transcriptionally inactive chromatin supporting the notion that ADAM10 ICD is actively involved in the suppression of stemness genes, such as *Krt19* and *Cd133*. The presence of nuclear ADAM10 ICD correlated with histone H3 trimethylation at Lys-27 (H3K27me3), which has been linked to the histone methyltransferase activity of polycomb repressive complex (PRC) 2 [39]. Interestingly, *Ezh1*^-/-^*Ezh2*^Δhep^ mice, deficient for enhancer of zeste homolog (EZH) 1 and 2, catalytic components of PRC2, displayed loss of H3K27me3 and an increase in A6^+^ HPCs which correlated with increased expression of *Cd24*, *Krt19* and *Cd133* [40]. It is therefore tempting to speculate, that ADAM10 ICD is actively involved in the recruitment of PRC2 to stem cell genes. We further demonstrate that in HPCs ADAM10 is proteolytically processed by ADAM9 and to a lesser extent by ADAM15. Consistently, *Adam9* and *Adam15* expression was elevated in human and murine livers under chronic liver disease conditions and nuclear ADAM10 was detected in HPCs and hepatocytes in patients with chronic liver disease. Furthermore, nuclear translocation of the hepatic transcription factor HNF4α was linked to ADAM10 ICD nuclear localization, suggesting an essential function of ADAM10 ICD in HNF4α nuclear shuttling. Therefore, a complex of ADAM10 ICD and HNF4α may be responsible for the transcriptional suppression of HPC stemness genes, shifting HPCs into a differentiation-permissive state (Fig.S9a+b). Interestingly, a recent report demonstrates the importance of protein interactions with the ADAM10 C-terminus for pre-synaptic plasticity [41]. Therefore, the ADAM10 C-terminus appears to be involved in multiple protein-protein interactions confering non-catalytic functions of ADAM10.

Taken together, our novel data demonstrate that proteolytic processing of ADAM10 by ADAM9 but not proteolytic activity of ADAM10 itself in HPCs is essential for the differentiation of HPCs. The loss of ADAM10 inhibits HPC differentiation and has therefore been demonstrated to play a critical role in aggravating chronic liver disease. For the first time, we assign a functional role to an ADAM protease soluble intracellular domain. Transcriptional and potentially epigenetic regulation by ADAM10 ICD in HPCs is a completely novel concept that might expand to other adult tissue stem cells.

## Acknowledgment

We thank Fabian Neumann and Pit Christoffersen for excellent technical assistance. We are grateful to the members of the Victor-Hensen animal facility at the Christian-Albrechts-University Kiel and the animal facility staff at Curtin University in Perth.

This work was supported by the Deutsche Forschungsgemeinschaft (DFG), Bonn [grant number SFB841; Liver inflammation: Infection, immune regulation and consequences, to D.S.-A., S.R.-J., C.S. and J.H.; grant number SFB877 to S.R.-J., P.S.; KFO306 to C.S. and J.H.], the Australian Technology Network/Deutscher Akademischer Austauschdienst [ATN-DAAD PPP to D.S.-A. and J.T.-P.], the Cluster of Excellence ‘Inflammation at Interfaces’ to S.R.-J. and the National Health and Medical Research Council of Australia (Grant numbers APP1061332 to G.A.R., and APP1087125 and APP1160323 to G.A.R, J.T.-P and J.K.O.).

## Author contributions

B.W. performed most of the experiments and analyzed data, M.M., J. G-T., J.B. and N.S. performed animal experiments and contributed to writing the manuscript. M.P. performed meta-analysis of human and mouse transcriptome data, L.G., R.P., S.W. performed experiments and analyzed data. S.R-J., P.S. contributed to writing the manuscript. R.B. provided Mdr2^-/-^ samples. D.S., C.S, C.J., L.S., J.K.-G., J.K.O. and J.H. provided and analyzed human samples. J.T.-P., G.A.R. provided material, analyzed data and contributed to writing of the manuscript. D.S.-A. conceived the study, supervised experiments, analyzed data and wrote the manuscript.

