## Supplementary Material for "A non-catalytic function of a disintegrin and metalloprotease 10 determines hepatic progenitor cell fate"

### Supplementary Experimental Procedures

*Cell lines*

HPC lines BMOL[1] (murine) and HepaRG[2] (human) were previously described. BMOL cells were cultured in William's E medium, supplemented with 2% FBS, 2 mM L-glutamine, 15 ng/ml rh insulin-like growth factor (IGF) II, 10 ng/ml rmEGF and 5 µg/ml rhInsulin. HepaRG cells were cultured in William's E medium supplemented with 10% FCS, 5 µg/ml rhInsulin and 50 µM hydrocortisone hemisuccinate. All cells were cultured at 37°C, 5 % $CO_2$ and 95 % relative humidity (RH).

*Generation of BMOL cells with insertion of 3xFLAG cassette into the Adam10 locus*

sgRNAs targeting exon 16 of murine Adam10 were cloned into pLentiCRISPRv2 as previously described[3]. *Adam10* 5’ (HOM1) and 3’ (HOM2) homology regions were PCR-amplified from BMOL genomic DNA. The Primer sequences are available on request. Homology arms and the 3xFLAG-2A-Neomycin cassette from the pFETCH plasmid were fused by Gibson assembly as previously described[4]. BMOL cells were transfected with sgRNA constructs and repair template using Lipofectamine 3000 and transfected cells were selected with 0.25 mg/ml G418.

*siRNA transfection and inhibitor treatment*

Cells were seeded in a 6-well plate at a final concentration of 2.5 x$10^5$ cells per well. Cells were subsequently transfected with 48 pmol siRNA/well using Interferrin (Polyplus transfection) transfection reagent, according to the manufacturer's instructions. Medium

was replaced and siRNA transfection was repeated after 48 h. Cells were then cultured for another 24 h for the final experiment.
Cells were treated with 100 nM SR9243, 3 µM GI254023X or 100 ng/ml recombinant tissue inhibitor of metalloproteinase (TIMP)-1 4 h after cell seeding and prior to siRNA transfection and replaced with fresh inhibitor solution every 24 h.
For broad metalloprotease inhibition, BMOL cells were treated with 10 µM Marimastat for 5 days with replacement of inhibitor-containing medium every 24h.

*Tube formation assay*

A 24-well plate was coated with 300 µl of Matrigel basement matrix (Corning, protein conc. > 10 mg/ml). $1x10^5$ cells/ml were seeded on top of the Matrigel in 300 µl/well. If indicated, siRNA treatment was performed 72 h prior to seeding. DAPT was added to the cell suspension at a final concentration of 2 µM. After incubation for 24 h under normal cell culture conditions, images of formed tubes were taken with an AZ100 microscope with a DS-Fi2 camera (Nikon).

*Immunohistochemistry and immunofluorescence*

Liver tissues were fixed in phosphate buffered saline (PBS) containing 4% formaldehyde, dehydrated and embedded in paraffin, and cut into 4 µm-thick sections. Sections were stained with haematoxylin-eosin or Sirius Red using standard protocols. Additional sections were stained immunohistochemically. Primary antibodies were detected by biotinylated secondary antibodies and subsequently stained using a peroxidase DAB kit (Dako, Hamburg, Germany). Antibody specifications are listed in Supplementary Table 3. Haematoxylin was used for counterstaining. Alternatively, murine tissue specimens were snap frozen in OCT TissueTek compound (Plano GmbH, Wetzlar, Germany) and cut into 8 µm-thick sections. Frozen human tissue specimens were embedded in OCT TissueTek compound and cut into 8 µm-thick sections. Sections were fixed in acetone:methanol (1:1 v/v) for 2 min and subjected to immunofluorescent staining according to standard procedures. Antibody specifications are listed in Supplementary Table 3.
For immunofluorescent studies, cells were grown on cover slips and fixed in 4% paraformaldehyd in PBS. Cells were permeabilized in 0.2% saponin in PBS and quenched in 0.12% glycin and 0.2% saponin in PBS. Cells were incubated with primary antibodies (Supplementary Table 2) overnight at 4°C and detected with Alexa Fluor® 488, Alexa Fluor® 594 or Alexa Fluor® 647 labeled secondary antibodies (1:200, Life Technologies, Darmstadt, Germany) for 1 hour at room temperature.

Coverslips were mounted with MOWIOL/DABCO with 1 µg/ml DAPI (Life Technologies, Darmstadt, Germany). Cells were then analysed on an Olympus FluoView 1000 confocal microscope.

*Immunoblotting*

Samples were lysed in RIPA buffer supplied with 50 mM NaF, protease and phosphatase inhibitors. Proteins were separated by electrophoresis on 10 % SDS gels and transferred to PVDF or nitrocellulose membranes. Membranes were incubated with primary antibodies (Supplementary Table 2) overnight at 4°C and with horseradish peroxidase-conjugated secondary antibodies at room temperature for 1h. An ECL substrate kit was used for detection (Thermo Scientific, St. Leon-Rot, Germany).

*Quantitative real time PCR*

Total RNA was isolated from cryopreserved whole liver tissue using TRIzol (LifeTechnologies, Darmstadt, Germany) according to the manufacturer's instructions. One microgram of total RNA was used for reverse transcription with oligo-(dT)18 primers and RevertAid Reverse Transcriptase (Thermo Scientific, St. Leon-Rot, Germany) and subsequently subjected to quantitative polymerase chain reaction. Respective gene expression levels were normalised to the expression of β-actin or tubulin and expressed as fold changes relative to normalised levels in the corresponding control mice. Primer sequences are detailed in Supplementary Table 1.

*Gene expression metaanalysis*

The normalised microarray gene expression data of GSE28619[5] and GSE33161[6] were downloaded from the Gene Expression Omnibus database. The GEO2R online tool (https://www.ncbi.nlm.nih.gov/geo/geo2r/) with Benjamini & Hochberg multiple testing correction[7] was used to analyse differentially expressed genes. The expression of ADAM9, 10, and 15 were selected from differentially expressed genes.

### Supplementary Figures

##### Supplementary Figure 1

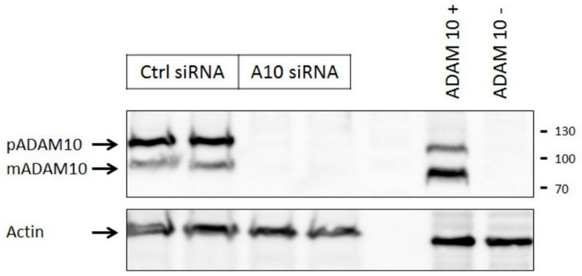


**Supplementary Figure 1.** siRNA-mediated suppression of ADAM10 expression in BMOL cells was verified by SDS-PAGE and immunoblotting. pADAM10: ADAM10 pro-form; mADAM10: ADAM10 mature form.

**Supplementary Figure 2**

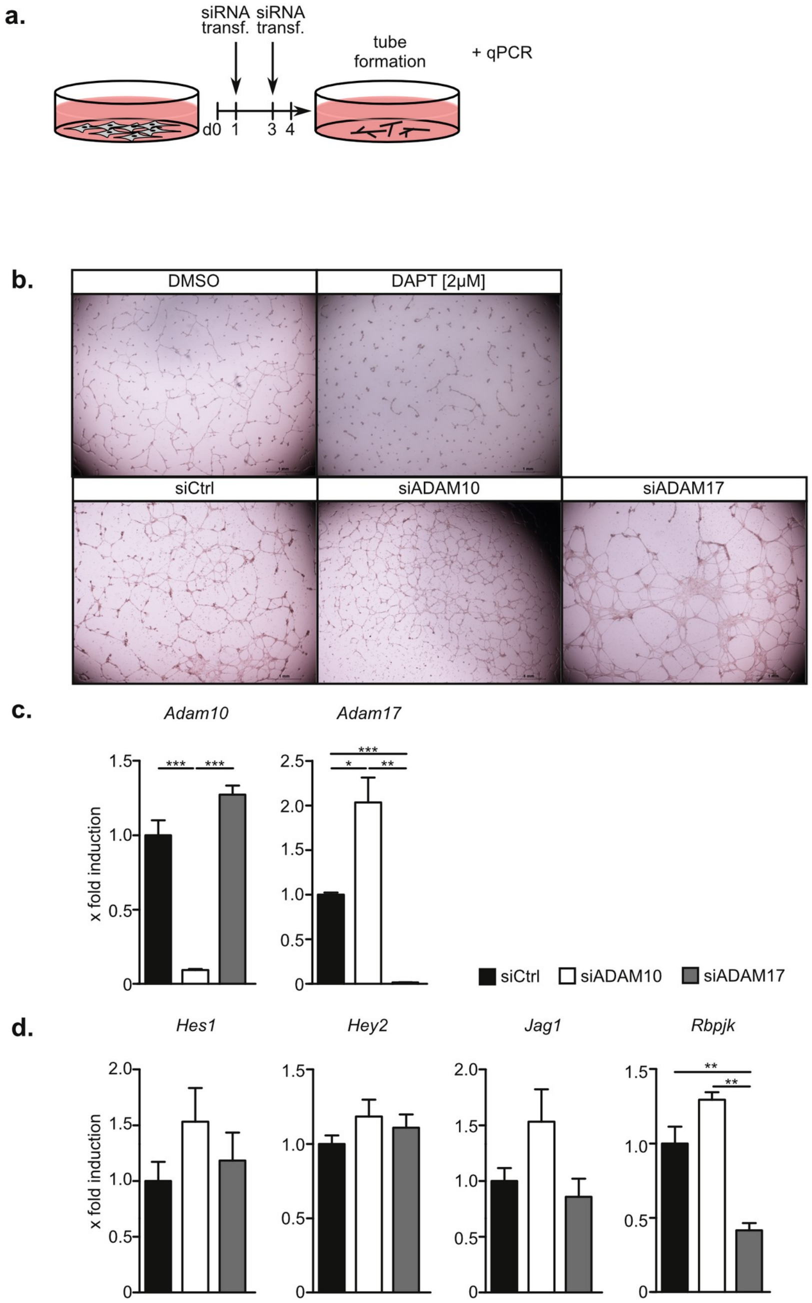


**Supplementary Figure 2. ADAM10 is not required for Notch signaling in HPCs. a.** Experimental outline as performed in b-d. **b.** Notch2-dependent tubulogenesis of BMOL cells on matrigel is impaired under γ-secretase inhibition with DAPT but not in the absence of ADAM10. **c.** ADAM10 but not ADAM17 expression is significantly downregulated by siRNA transfection in BMOL cells. **d.** Expression of Notch target genes *Hes1* and *Hey2*, Notch-ligand *Jag1* and co-regulator *Rbpjk* is not changed in the absence of ADAM10. Expression of the indicated genes was assessed by qRT-PCR. Data are mean ± s.e.m. n=3 (c,d) *$P<0.05$, **$P<0.01$, ***$P<0.001$, Two-tailed unpaired Student's t-test (c,d).

### Supplementary Figure 3

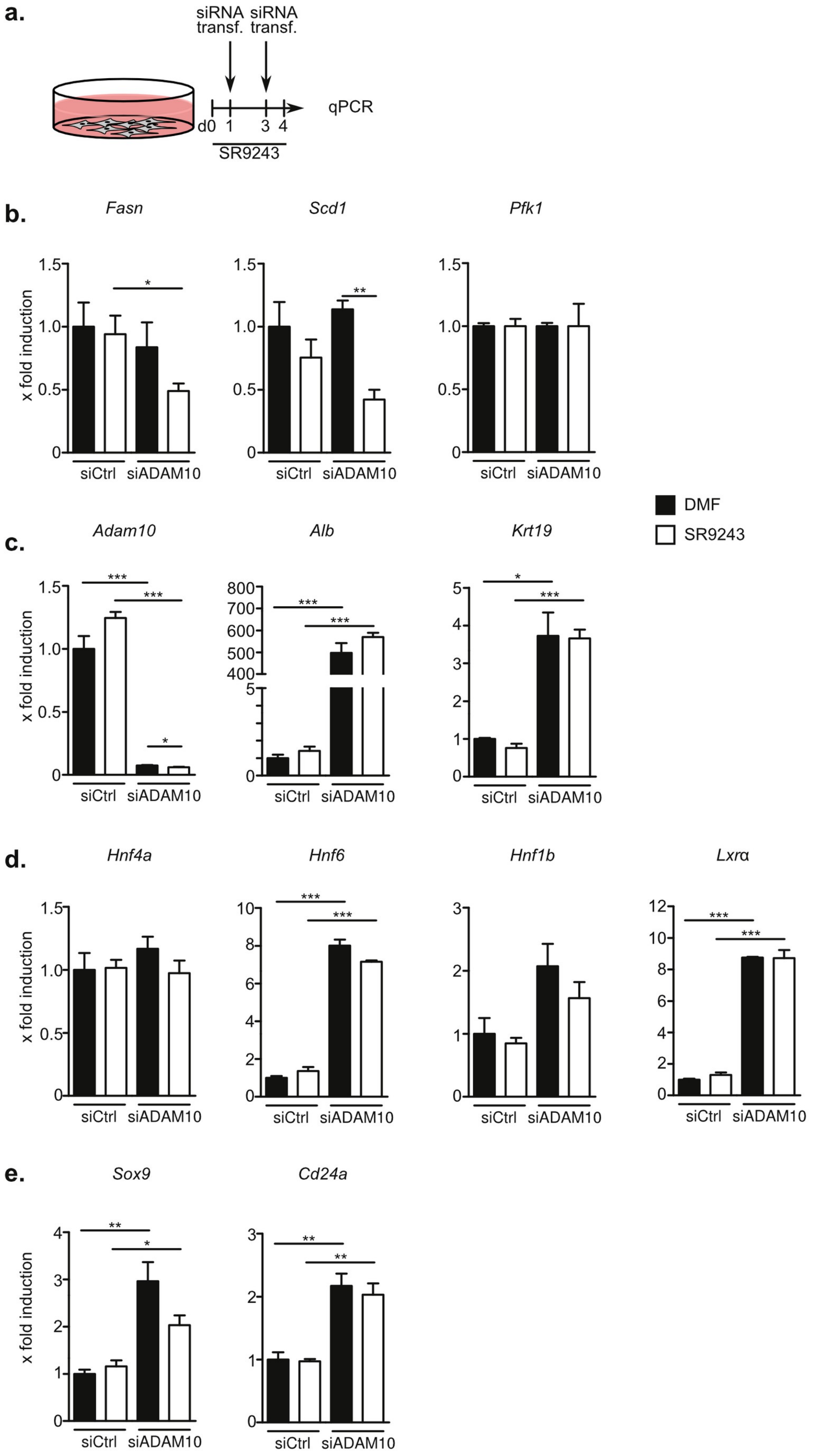

**Supplementary Figure 3: Transcription factor LxRα is dispensable for the induction of stemness-related gene expression in ADAM10-deficient HPCs. a.** Experimental setup for experiments performed in b-e. **b.** Expression of the LxRα target gene Scd1 is downregulated in the presence of the inhibitory inverse LxRα agonist SR9243. **c.** Induction of albumin and *Krt19* expression in ADAM10-deficient HPCs is not impaired in the presence of LxRα inhibition. **d.** LxR does not regulate hepatic transcription factors in ADAM10-deficient HPCs. **e.** Stemness-related genes *Sox9* and *Cd24a* are not regulated by LxRα in ADAM10-deficient HPCs. Data are mean ± s.e.m. n=3 (b-d) *$P$<0.05, **$P$<0.01, ***$P$<0.001, Unpaired, two-tailed Student's t test (b-e)

**Supplementary Figure 4**

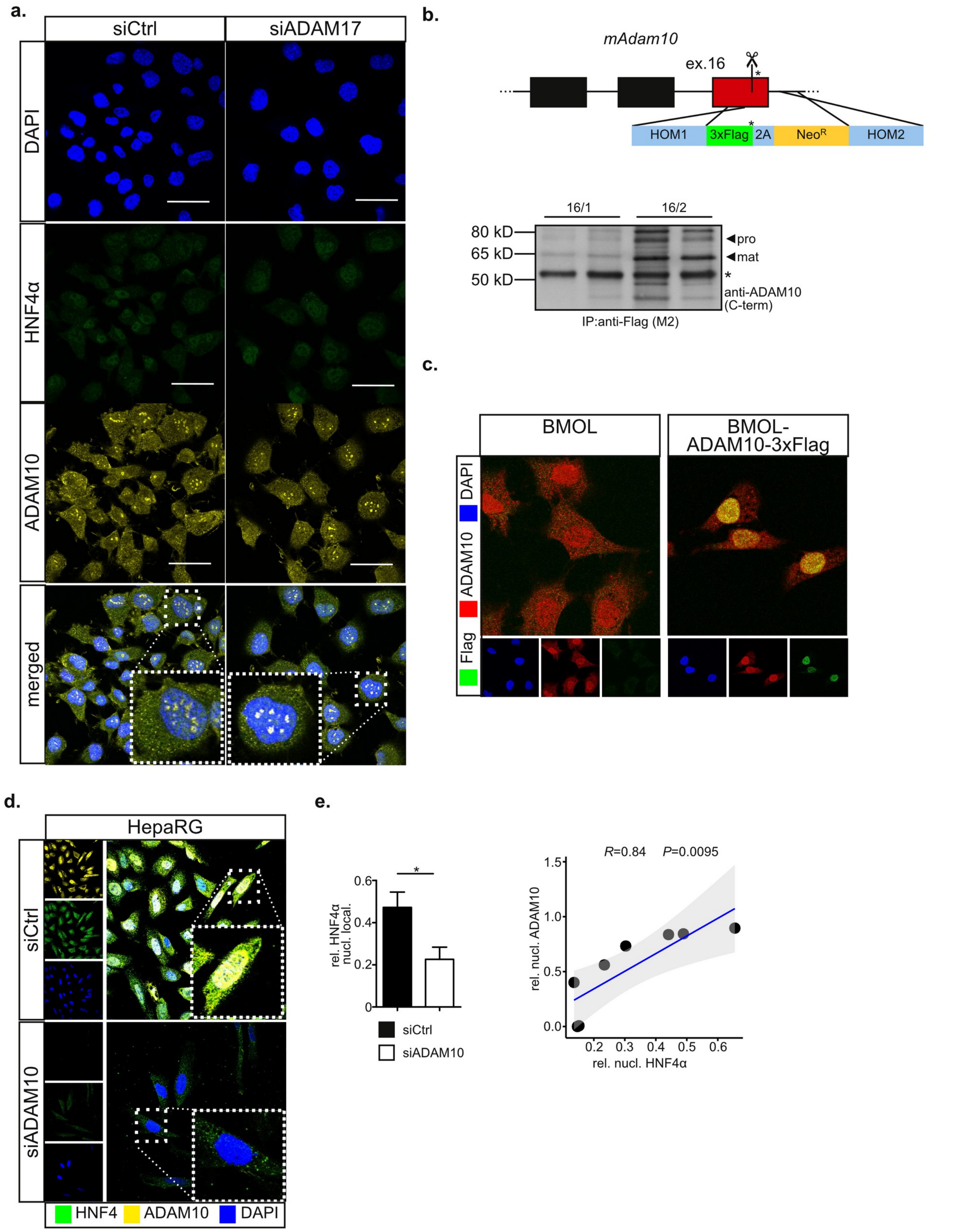


**Supplementary Figure 4. a.** Nuclear translocation of ADAM10 and HNF4α is not altered in the absence of ADAM17. **b.** upper panel: Targeting strategy for CRISPR-mediated endogenous Flag-tagging of the *Adam10* locus. sgRNAs 16/1 and 16/2 target exon 16 close to the endogenous Stop codon. The targeting construct contained two homology arms (HOM1, 2), the Flag-tag, a 2A site and a neomycin resitence cassette ($Neo^R$). lower panel: Flag-tagged ADAM10 was immunoprecipitated using anti-Flag antibodies and detected by immunoblotting using antibodies against ADAM10 C-terminus. **c.** Immunofluorescence analysis of BMOL-ADAM10-3xFlag confirms nuclear localisation of ADAM10 ICD using anti-Flag antibodies. **d.** HNF4α nuclear translocation is also dependent on ADAM10 in the human HPC line HepaRG. **e.** Nuclear translocation of HNF4α correlates with nuclear localisation of ADAM10 in the human HPC line HepaRG. Data are mean ± s.e.m. n=7 (c, from 2 independent experiments) **P*<0.05, Unpaired one-tailed Mann-Whitney U (e), Pearson correlation (e).

**Supplementary Figure 5**

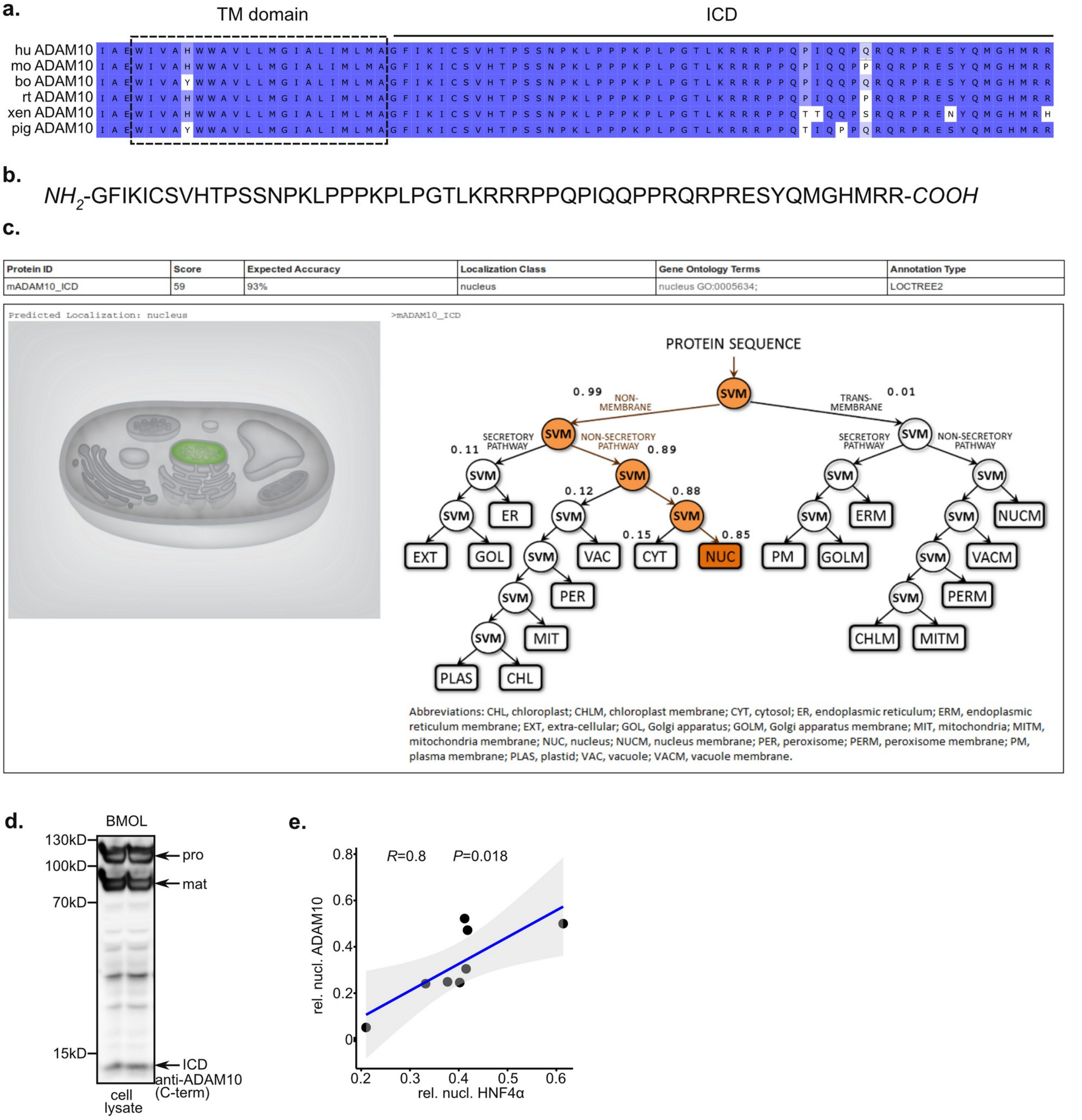


**Supplementary Figure 5. ADAM10 ICD sequence predicts nuclear localisation. a.** Multiple sequence alignment of the ADAM10 intracellular domain (ICD) reveals high conservation among the indicated species. **b.** Aminoacid sequence of murine ADAM10 ICD used for LocTree3 prediction. **c.** LocTree3 predicts nuclear localisation of ADAM10 ICD, based on multiple sequence alignments. **d.** Limited proteolysis generates an ADAM10 intracellular domain (ICD) in HPCs as assessed by immunoblotting using an antibody directed against the C-terminus of ADAM10. **e.** Nuclear localization of HNF4α significantly correlates (Pearson correlation) with nuclear localization of ADAM10 ICD in Marimastat-treated BMOL cells.

**Supplementary Figure 6**

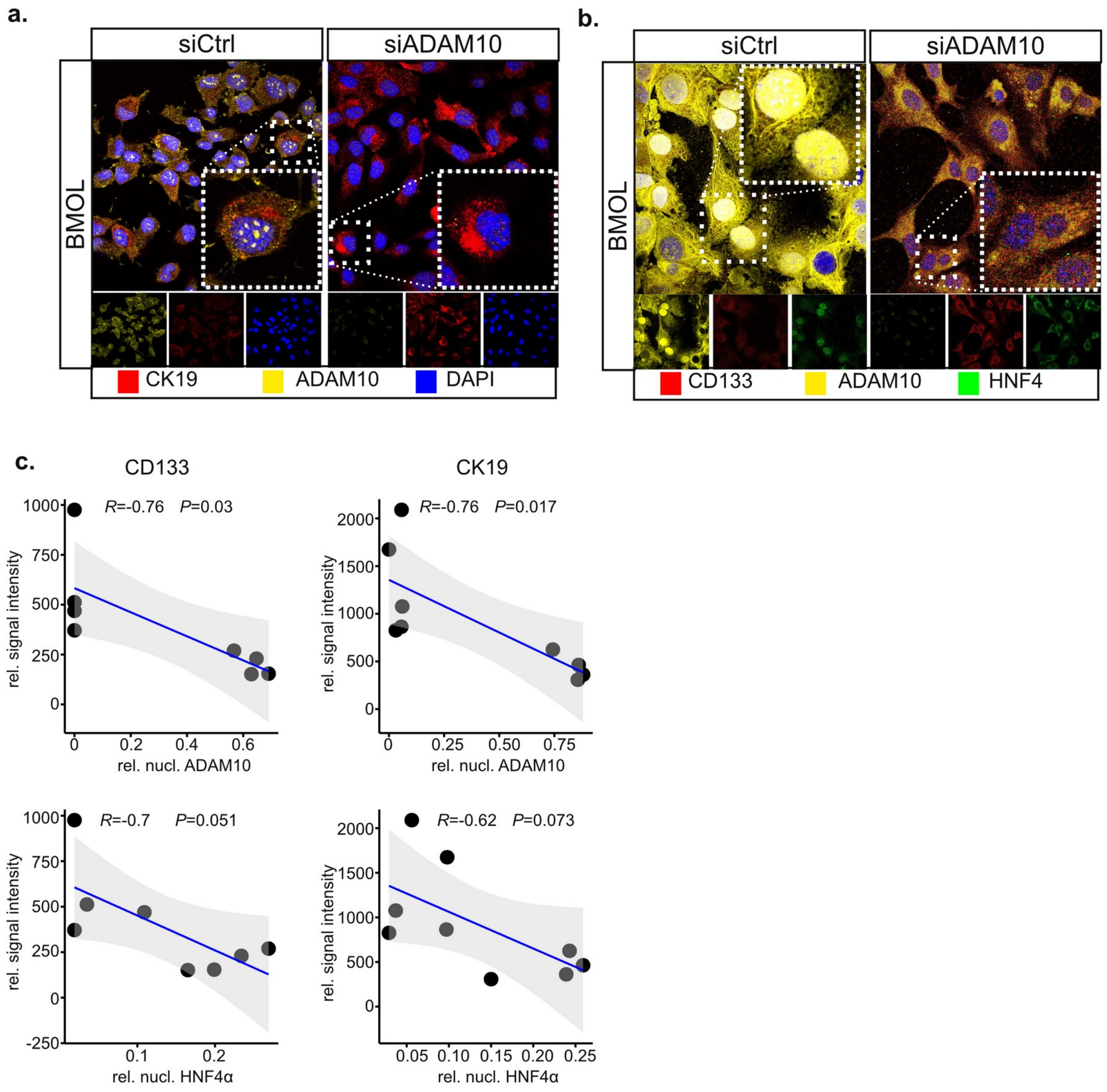


**Supplementary Figure 6. a-c.** Upregulation of CK19 and CD133 inversely correlates with nuclear translocation of ADAM10 and HNF4α as determined by immunofluorescent staining of siRNA-transfected BMOL cells with the indicated antibodies. n=8-9 (from 2-3 independent experiments), Pearson correlation (c).

**Supplementary Figure 7**

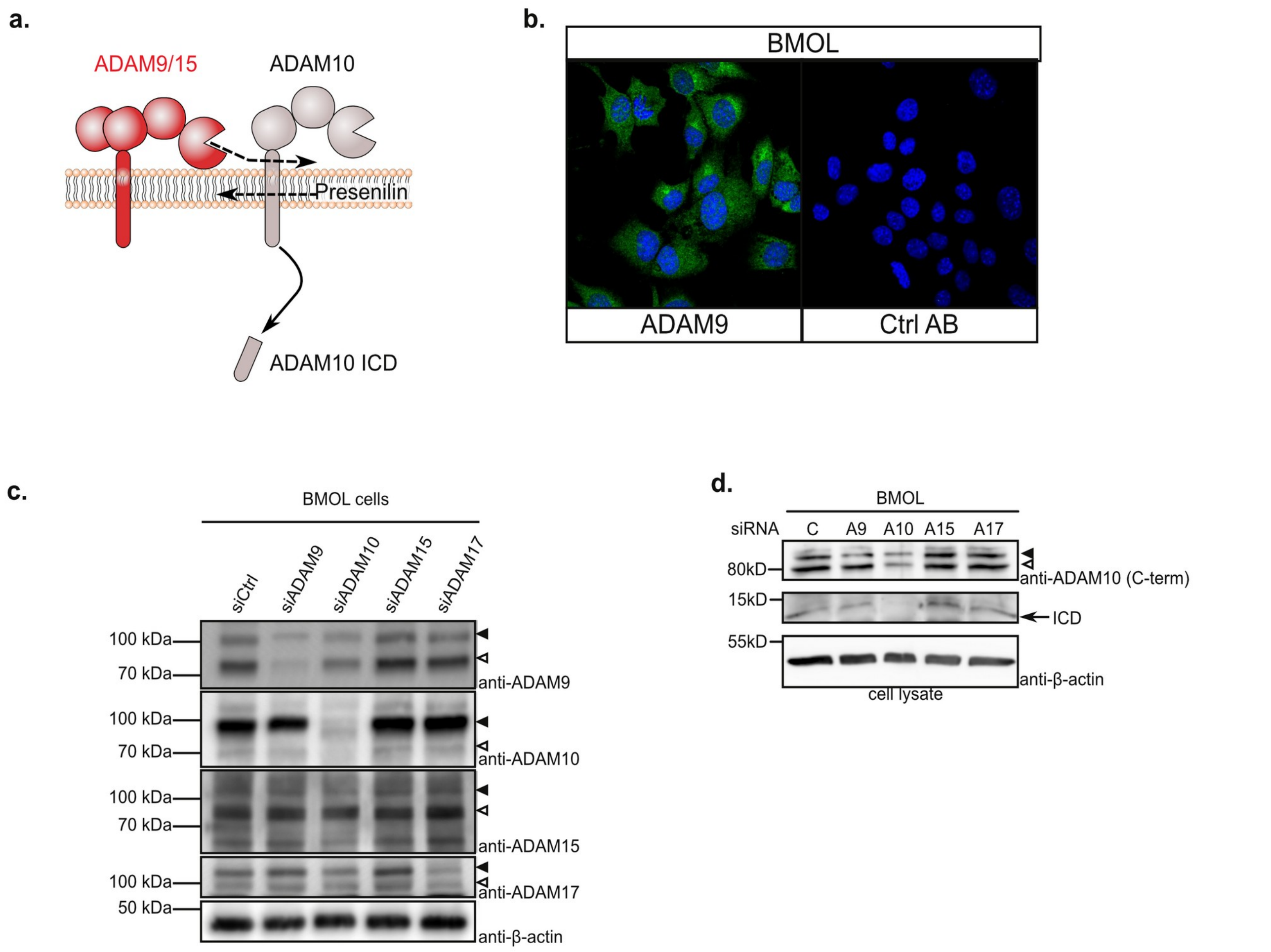


**Supplementary Figure 7. ADAM10 is processed by ADAM9 and 15 in HPCs. a.** ADAM10 can be proteolytically processed by ADAM9 or ADAM15. The γ-secretase complex with its catalytic component Presenilin subsequently generates an ADAM10 ICD. **b.** ADAM9 is expressed in the murine HPC line BMOL as assessed by immunofluorescence. **c.** Verification of siRNA-mediated suppression of the indicated ADAM proteases by immunoblotting. Filled arrow heads mark pro-form, open arrow heads mature form of the indicated ADAM proteases. **d.** ADAM10 ICD formation is impaired in the absence of ADAM9 as assessed by immunoblotting using an antibody directed against the C-terminus of ADAM10. Filled arrow heads mark pro-form, open arrow heads mature form of the indicated ADAM proteases.

**Supplementary Figure 8**

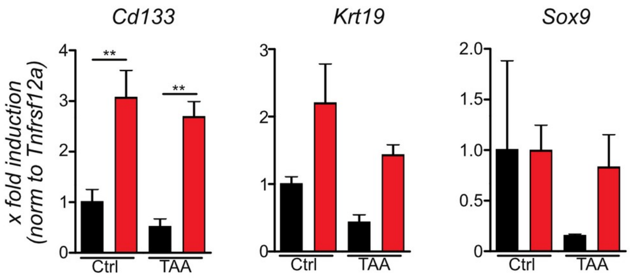


**Supplementary Figure 8. CD133 is increased in ADAM10-deficient HPCs.** Relative expression of the indicated genes was normalised to *Tnfrsf12a*-expression as an indicator of LPC number. Data are mean ± s.e.m. n=3 mice/group *$P$<0.05, one-way ANOVA on ranks with Bonferroni's post-hoc test

**Supplementary Figure 9**

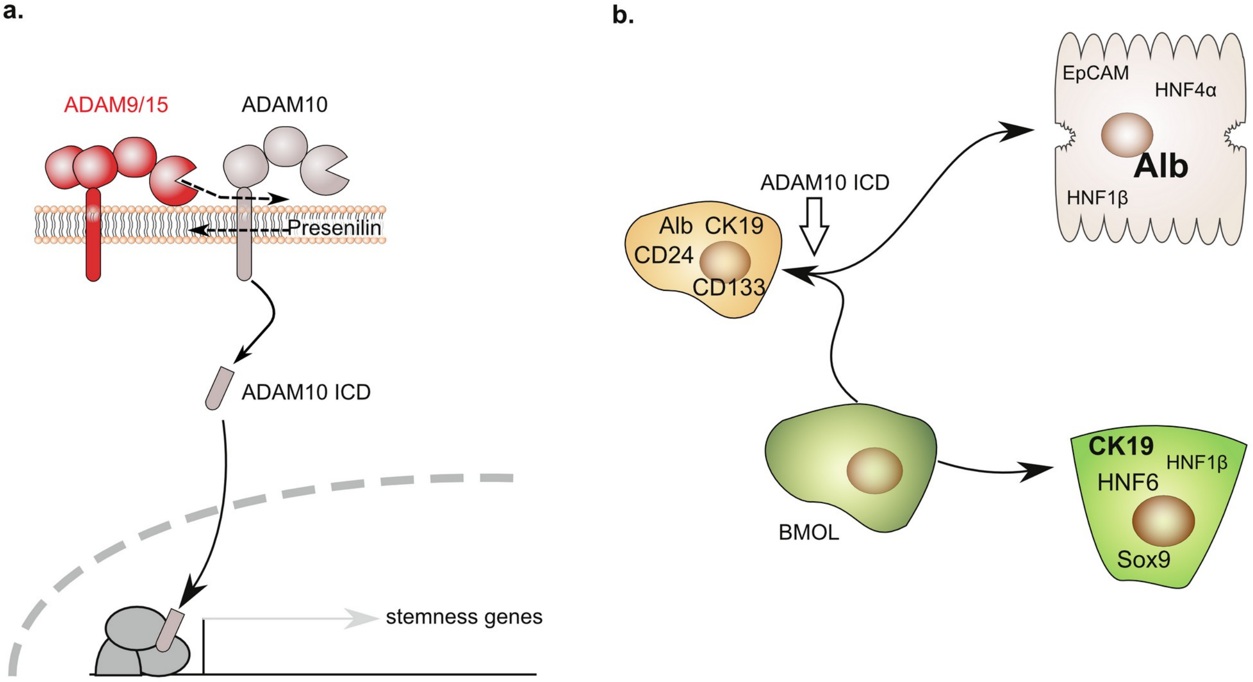


**Supplementary Figure 9. Proteolytic processing of ADAM10 is a prerequisite for HPC differentiation. a.** In HPCs, ADAM10 is proteolytically processed by ADAM9/15 and presumably the Presenilin-containing γ-secretase complex to generate an ADAM10 intracellular domain (ICD). ADAM10 ICD translocates to the nucleus in order to suppress expression of genes associated with stemness. Furthermore, ADAM10 ICD might assist in HNF4α nuclear translocation. **b.** Nuclear translocation of ADAM10 ICD is a prerequisite for the differentiation of HPCs.

#### Supplementary tables

**Supplementary table 1** - siRNA sequences used in this study

| Target | target species | Distributed by | Product nr. |
|---|---|---|---|
| siCtrl | human/mouse | Dharmacon | D-001810-10-05<br>OnTarget Plus<br>Non-targeting |
| siADAM9-1 | mouse | ThermoScientific | MSS201740 |
| siADAM9-2 | mouse | ThermoScientific | MSS201741 |
| siADAM9-3 | mouse | ThermoScientific | MSS201742 |
| siADAM10-1 | mouse | ThermoScientific | MSS201701 |
| siADAM10-2 | mouse | ThermoScientific | MSS201702 |
| siADAM10-3 | mouse | ThermoScientific | MSS201703 |
| siADAM10 | human | Dharmacon | L-004503-00-0005 |
| siADAM15 | mouse | Dharmacon | L-057378-01 |
| siADAM17 | mouse | Dharmacon | L-040408-00 |

**Supplementary table 2** - qRT-PCR sequences used in this study

| Target gene | Forward Primer | Reverse Primer | UPL probe | Sequence Reference |
|---|---|---|---|---|
| *Adam9* | 5’ TTTCTCCGGCAGTGAGTACA 3’ | 5’ GCATTGAAGCTTTCCACACA 3’ | 48 | NM_007404.2<br>NM_001270996.1<br>XM_006508986.3 |
| *Adam10* | 5’ GGGAAGAAATGCAAGCTGAA 3’ | 5’ CTGTACAGCAGGGTCCTTGAC 3’ | 38 | NM_007399.4 |
| *Adam15* | 5’ AGCACAGGAATGTCGAAGAAA 3’ | 5’ TTGAGCTGGGTCATGCAGT 3’ | 104 | NM_009614.3<br>NM_001037722.3<br>XM_006500918.1<br>XR_375482.3 |
| *Adam17* | 5’ TGTGGTTATTATTTAAATGCAGATAGTGA 3’ | 5’ TCACTCGACGAACAAACTCTTC 3’ | 38 | NM_009615.6<br>NM_001277266.1<br>NM_001291871.1 |
| *Alb* | 5’ TGACCCAGTGTTGTGCAGAG 3’ | 5’ TTCTCCTTCACACCATCAAGC 3’ | 1 | NM_009654.4 |
| *Hnf4a* | 5’ CAGCAATGGACAGATGTGTGA 3’ | 5’ TGGTGATGGCTGTGGAGTC 3’ | 27 | NM_008261.3 |
| *LXR* | 5’ CGCGACAGTTTTGGTAGAGG 3’ | 5’ CTCCAGCCACAAGGACATC 3’ | 1 | AF085745.1 |

| | | | | |
|---|---|---|---|---|
| *Hnf1b* | 5’ GACACTCCTCCCATCCTCAA 3’ | 5’ CATGTATCCCTTGATCATTTTGG 3’ | 1 | |
| *Hnf6* | 5’ AGACCTTCCGGAGGATGTG 3’ | 5’ TTGCTCTTTCCGTTTGCAG 3’ | 92 | |
| *Krt19* | 5’ AGTCCCAGCTCAGCATGAA 3’ | 5’ TAACGGGCCTCCGTCTCT 3’ | 97 | NM_008471.3<br>NM_001313963.1 |
| *Sox9* | 5’ GTACCCGCATCTGCACAAC 3’ | 5’ CTCCTCCACGAAGGGTCTCT 3’ | 66 | AF421878.1 |
| *Cd133* | 5’ CTGCGATAGCATCAGACCAA 3’ | 5’ TATCCACTGATGGGAGCTGA 3’ | 32 | |
| *Cd24a* | 5’ CTTCTGGCACTGCTCCTACC 3’ | 5’ TGGTGGTAGCGTTACTTGGA 3’ | 38 | NM_009846.2 |
| *Cd44* | 5’ CTCCTTCTTTATCCGGAGCAC 3’ | 5’ TGGCTTTTTGAGTGCACAGT 3’ | 49 | |
| *Tub* | 5’ CTGGAACCCACGGTCATC 3’ | 5’ GTGGCCACGAGCATAGTTATT 3’ | 88 | |
| *Notch1* | 5’ CTGGACCCCATGGACATC 3’ | 5’ AGGATGACTGCACACATTGC 3’ | 80 | |
| *Notch2* | 5’ TGCCTGTTTGACAACTTTGAGT 3’ | 5’ GTGGTCTGCACAGTATTTGTCAT 3’ | 6 | |
| *Rbpjk* | 5’ AGTCTTACGGAAARGAAAAACGA 3’ | 5’ CCAACCACTGCCCATAAGAT 3’ | 63 | |
| *Jag1* | 5’ TGGCCGAGGTCCTACACTT 3’ | 5’ GCCTTTTCAATTATGCTATCAGG 3’ | 22 | |
| *Hes1* | 5’ TGCCAGCATGATATAATGGAGAA 3’ | 5’ CCATGATAGGCTTTGATGACTTT 3’ | 20 | |
| *Hey1* | 5’ CATGAAGAGAGCTCACCCAGA 3’ | 5’ GAACACAGAGCCGAACTCAA 3’ | 17 | |
| *Hey2* | 5’ GTGGGGAGCGAGAACAATTA 3’ | 5’ GTTGTCGGTGAATTGGACCT 3’ | 104 | |
| *Scd1* | 5’ CGTGATGTTCCAGAGGAGGTA 3’ | 5’ CGCAAGAAGGTGCTAACGA 3’ | 3 | |
| *Fasn1* | 5’ CAACATGGGACACCCTGAG 3’ | 5’ GTTGTGGAAGTGCAGGTTAGG 3’ | 1 | |
| *Pfk1* | 5’ GGAACTCAAGGGGAACTCG 3’ | 5’ AGCACAGTAGACCCCCACAT 3’ | 102 | |
| *Gapdh* | LifeTechnologies TaqMan Gene Expression Assays | | | NM_008084.2 |

**Supplementary table 3** - antibodies used in this study

| Target | Host species | Dilution | Distributed by | Product nr. |
|---|---|---|---|---|
| Immunoblot | | | | |
| ADAM10 (C-terminus) | rabbit | 1:1,000 | Millipore | 2382103 |
| ADAM9 | rabbit | 1:2,000 | GeneTex | 130081 |
| ADAM15 (ectodomain) | goat | 1:2,500 | R&D Systems | FHE021704A |
| ADAM17 | rabbit | 1:2,000 | Abcam | ab39162 |
| β-actin | mouse | 1:10,000 | Sigma-Aldrich | A1978 |
| H3 | rabbit | 1:2,000 | Cell Signaling | 4499 |
| c-Met | mouse | 1:1,000 | Cell Signaling | 3127 |
| SP1 | rabbit | 1:1,000 | Millipore | ABE135 |
| Immunofluorescence | | | | |
| ADAM10 (C-terminus) | rabbit | 1:100 | Millipore | 2382103 |
| HNF4α | goat | 1:100 | Santa Cruz | sc-6556 |
| CK19 | rat | 1:200 | DHSB | Troma-III |
| αSMA | rat | 1:200 | Dako/Agilent | |
| A6 | rat | 1:200 | Valentina Factor | |
| F4/80 | rat | 1:400 | BioRad | |
| ADAM9 | rabbit | 1:200 | GeneTex | 130081 |
| CD133 | rat | 1:200 | eBioscience | E04032-1632 |

| | | | | |
|---|---|---|---|---|
| H3 K9ac | mouse | 1:100 | ActiveMotif | 61252 |
| H3 K27me2m3 | mouse | 1:100 | ActiveMotif | 39537 |
| Immunohistochemistry | | | | |
| F4/80 | rat | 1:200 | Biorad | MCA497R |
| Gr1/Ly6G | rat | 1:400 | eBioscience | 14-5931 |
